# One to rule them all? Assessing the performance of sustainable forest management indicators against multitaxonomic data for biodiversity conservation

**DOI:** 10.1101/2024.02.12.579875

**Authors:** Yoan Paillet, Livia Zapponi, Peter Schall, Jean-Matthieu Monnet, Christian Ammer, Lorenzo Balducci, Steffen Boch, Gediminas Brazaitis, Alessandro Campanaro, Francesco Chianucci, Inken Doerfler, Markus Fischer, Marion Gosselin, Martin M. Gossner, Jacob Heilmann-Clausen, Jeňýk Hofmeister, Jan Hošek, Kirsten Jung, Sebastian Kepfer-Rojas, Peter Odor, Flóra Tinya, Giovanni Trentanovi, Giorgio Vacchiano, Kris Vandekerkhove, Wolfgang W. Weisser, Michael Wohlwend, Sabina Burrascano

## Abstract

Several regional initiatives and reporting efforts assess the state of forest biodiversity through broad-scale indicators based on data from national forest inventories. Although valuable, these indicators are essentially indirect and evaluate habitat quantity and quality rather than biodiversity *per se*. Therefore, their link to biodiversity may be weak, which decreases their usefulness for decision-making.

For several decades, Forest Europe indicators assessed the state of European forests, in particular their biodiversity. However, no extensive study has been conducted to date to assess their performance – i.e. the capacity of the indicators to reflect variations in biodiversity – against multitaxonomic data. We hypothesized that no single biodiversity indicator from Forest Europe can represent overall forest biodiversity, but that several indicators would reflect habitat quality for at least some taxa in a comprehensive way. We tested the set of Forest Europe’s indicators against the species richness of six taxonomic and functional groups across several hundreds of sampling units over Europe. We showed that, while some indicators perform relatively well across groups (e.g. deadwood volume), no single indicator represented all biodiversity at once, and that a combination of several indicators performed better.

Forest Europe indicators were chosen for their availability and ease of understanding for most people. However, we showed that gaps in the monitoring framework persist, and that surveying certain taxa along with stand structure is necessary to support policymaking and tackle forest biodiversity loss at the large scale. Adding context (e.g. forest type) may also contribute to increase the performance of biodiversity indicators.

**Impact statement:** Biodiversity indicators used to assess the state of Europe’s forests perform unequally; a combination of indicators is more informative

## 1. Introduction

Monitoring biodiversity in the face of global change is a challenge in many ecosystems across the world (IPBES 2019; Lindenmayer and Likens 2010). Despite large-scale initiatives such as GEOBON (Group on Earth Observation Biodiversity Observation Network 2008) and collaborative networks (Burrascano et al. 2023), no standard biodiversity monitoring scheme has been agreed in Europe or elsewhere. Long-term biodiversity monitoring hence currently relies on a limited number of initiatives used to assess the impact of climate change (e.g. Jiguet et al. 2012), habitat loss (Betts et al. 2017) or pollution (Rigal et al. 2023). Frameworks combining monitoring of biodiversity, types of pressure and ecosystem-level variables are rare, and may concern only single pressure vs. single taxon assessment (e.g. Proença et al. 2017; Weber et al. 2004), or use different methods for overall assessments (European Environment 2020). However, different taxonomic and functional groups likely respond differently to a given pressure, which challenges prioritization of policy initiatives and tools (Pereira and David Cooper 2006).

Forest ecosystems are no exception to the rule, and lack broad-scale standardized biodiversity and pressure data assessment. While forestry remains one of the main pressures that affect forest habitats in Europe (European Environment 2020), sustainable forest management is one of the main approaches to limit its impact (European Commission 2021). A series of agreed criteria and indicators, applied at several spatial scales (Forest Europe 2015) are used to tackle the different components of sustainable forest management, to develop novel management methods and best practices including biodiversity maintenance, conservation and enhancement of biodiversity.

Indicators are measurable parameters that allow to monitor the status and changes of forests in quantitative, qualitative and descriptive terms in relation to each criterion (FAO 2024). In the sustainable forest management framework, indicators are developed accounting for: i. a review of existing forest information; ii. ‘Bottom-up’ stakeholder engagement, i.e., priorities for forest sector performance; iii. ‘Top down’ alignment to international Criteria and Indicators (Linser and O’Hara 2019).

Given this basis, it comes as no surprise that National Forest Inventories were the first choice to develop nationwide indicators of sustainable forest management (Simons et al. 2021), since they are widely available and were originally designed to be representative of forests at a national scale. At the European level, data from National Forest Inventories have been aggregated and published every five years for more than three decades (Forest Europe 2020) and provide a set of indicators to inform sustainable forest management. For European forests, the Criterion 4 “*Maintenance, conservation and appropriate enhancement of biological diversity in forest ecosystems*” is the one that should help decision makers to assess progress towards biodiversity-friendly forest management.

However, the desired indication on biodiversity sustainability is not ensured, since most of the indicators are indirect (structural) proxies derived from National Forest Inventories or land cover maps (e.g. deadwood or forest fragmentation), while only two indicators involve other species than trees (namely 4.8 Threatened forest species, 4.10 Common forest bird species, Table 1). This approach follows the general trend of conservation programs and research to rely heavily on habitat attributes and area-based metrics, mainly in relation to the relative ease with which these measures can be collected and analyzed, compared with complex ecological or population processes (Marshall et al. 2020). As a matter of fact, despite the large use of National Forest Inventories data to indirectly assess the state and evolution of biodiversity (Chirici et al. 2012; Heym et al. 2021; Reise et al. 2019), these were often found to lack links with species indices (Gao et al. 2015; Paillet et al. 2018; Storch et al. 2023; Zeller et al. 2022). Despite a large corpus of individual studies and few syntheses (Zeller et al. 2023), no global and systematic assessment of the correlations between multi-taxonomic biodiversity and Forest Europe indicators has been attempted. This is particularly worrying since the European system of Criteria and Indicators is widely used as a reference across several regions, as well as for global initiatives (Linser and O’Hara 2019).

**Table 1:** Number of sampling units used to fit the models distributed per taxonomic and functional groups and Regeneration origin or Naturalness (Forest Europe 2020).

| Taxa | I4.2.Regeneration |  |  | I4.3.Naturalness |  |  | Total |  |
| --- | --- | --- | --- | --- | --- | --- | --- | --- |
|  | Coppice | Natural | Planting | Plantation | Semi-natural | Unmanaged | Sampling units | Sites |
| **Tracheo**<br>**phytes** | 10 | 734 | 94 | 94 | 550 | 194 | 838 | 43 |
| **Bryoph**<br>**ytes** | 3 | 353 | 66 | 66 | 198 | 158 | 422 | 30 |
| **Beetles** | 7 | 408 | 94 | 94 | 255 | 160 | 509 | 30 |
| **Birds** | 10 | 721 | 94 | 94 | 537 | 194 | 825 | 41 |
| **Fungi** | 8 | 502 | 94 | 94 | 347 | 163 | 604 | 36 |
| **Lichens** | 10 | 332 | 92 | 92 | 333 | 9 | 434 | 23 |

In this context, our goal was to provide an assessment of the performance of biodiversity indicators. Here, we define performance as the capacity of the indicator to reflect the variations 6 of one or more *indicanda* (defined as the parameter to be indicated, here an index of biodiversity). This encompasses the magnitude of the correlation between a given indicator and an *indicanda*, but also the respective fit of each indicator for a given *indicanda* as well as the relative fit of individual vs. combined indicators. We intended to support potential improvements in the global reporting processes for biodiversity conservation and sustainable forest management. In other words, we aimed to: i. assess the link between indicators and several *indicanda* (i.e. taxonomic and functional groups) based on a large dataset combining species and forest characteristic data; ii. identify the indicators that performed best and universally - i.e. for all groups; iii. define, if possible, the most effective combination of indicators for forest biodiversity.

To address these aims, we tested the performance of Forest Europe’s biodiversity indicators against six taxonomic and functional groups. We used a three step approach: we first tested individual indicators against the scaled species richness of the six groups to assess the magnitude of the links between single indicators and *indicanda*; second, we used a model selection approach to assess whether one single indicator outperformed the others; third, we tested whether several indicators performed better than single ones by searching for the best and most parsimonious combination of biodiversity indicators. We used a multi-taxonomic database (Burrascano et al. 2023; Chianucci et al. in press; Trentanovi et al. 2023) combining species inventories and forest structure to analyze the correlations between Forest Europe’s biodiversity indicators with the biodiversity of six groups (tracheophytes, epixylic and epiphytic bryophytes, birds, saproxylic beetles, saproxylic non-lichenized fungi and epixylic and epiphytic lichenized fungi).

## 2. Materials and Methods

### 2.1. Database

We used the database gathered within the framework of the COST action “BOTTOMS-UP” (CA18207 – Biodiversity Of Temperate forest Taxa Orienting Management Sustainability by Unifying Perspectives). In a nutshell, this unique database spatially combines sampling units 7 for both forest measurements and species of at least 3 taxa out of which at least one animal (Burrascano et al. 2023). It merges 34 different datasets from 12 European countries and more than 3500 sampling units. It covers a wide range of forest categories and environmental conditions but without apparent bias concerning the temperature and precipitation (Appendix 1, see Burrascano et al. 2023 for further details). It should be noted that temperate forest types are more represented than boreal ones (Burrascano et al. 2023).

The database is particularly well suited to test correlations between Forest Europe’s biodiversity indicators and biodiversity, since the former can be extracted either from forest measurement, species inventories as well as external (cartographic) data. From this database, we extracted the six most represented taxonomic and functional groups, namely tracheophytes, epixylic and epiphytic bryophytes (hereafter bryophytes), birds, saproxylic beetles, saproxylic non-lichenized fungi (hereafter fungi) and epixylic and epiphytic lichenized fungi (hereafter lichens).

While tracheophytes and birds were inventoried without any specific selection of the guilds targeted, only epiphytic and epixylic bryophytes and lichens were included (sampled on living trees and deadwood), and saproxylic fungi and beetles (dependent on deadwood substrates or on other organisms inhabiting deadwood).

Since the database is the result of the merging of different research projects, sampling designs and protocols varied across datasets (Burrascano et al. 2023; Trentanovi et al. 2023). Therefore, we scaled the species richness (number of species per sampling unit) by dividing it by the asymptotic gamma richness at the site level, with site representing a homogeneous sampling area with a maximum size of a few thousand hectares. The full dataset used in this study comprises between 23 and 43 sites depending on the taxa (Table 1). We derived site gamma diversity through sampling unit-based rarefaction curves using the R package iNEXT (Hsieh et al. 2016). Sites with less than 6 sampling units were discarded from the final database since the estimation of the gamma richness was judged non-reliable. Although species richness represents only a partial measure of biodiversity (Noss 1990), it was the simplest, widely used and most reliable metric that could be standardized for sampling effort. The effect of different standardization is presented in the Appendix 2 (Figure 2-1): since it did not show strongly different trends (Figure 2-2), we kept gamma-scaled richness as a way to control for heterogeneity in sampling efforts as well as regional species pools.

Exploratory analyses revealed especially large deadwood volumes linked with small sampling units (nugget effect). To avoid strong leveraging from these outliers (Zuur et al. 2010), we limited the maximum deadwood volume per sampling unit to 500 m^3^/ha, a value that corresponds to the maximal volumes observed in primeval forest of Europe (e.g. Christensen et al. 2005). The final data distribution per taxonomic and functional groups is shown in Table 1 and Figure 1.

**Figure 1:**
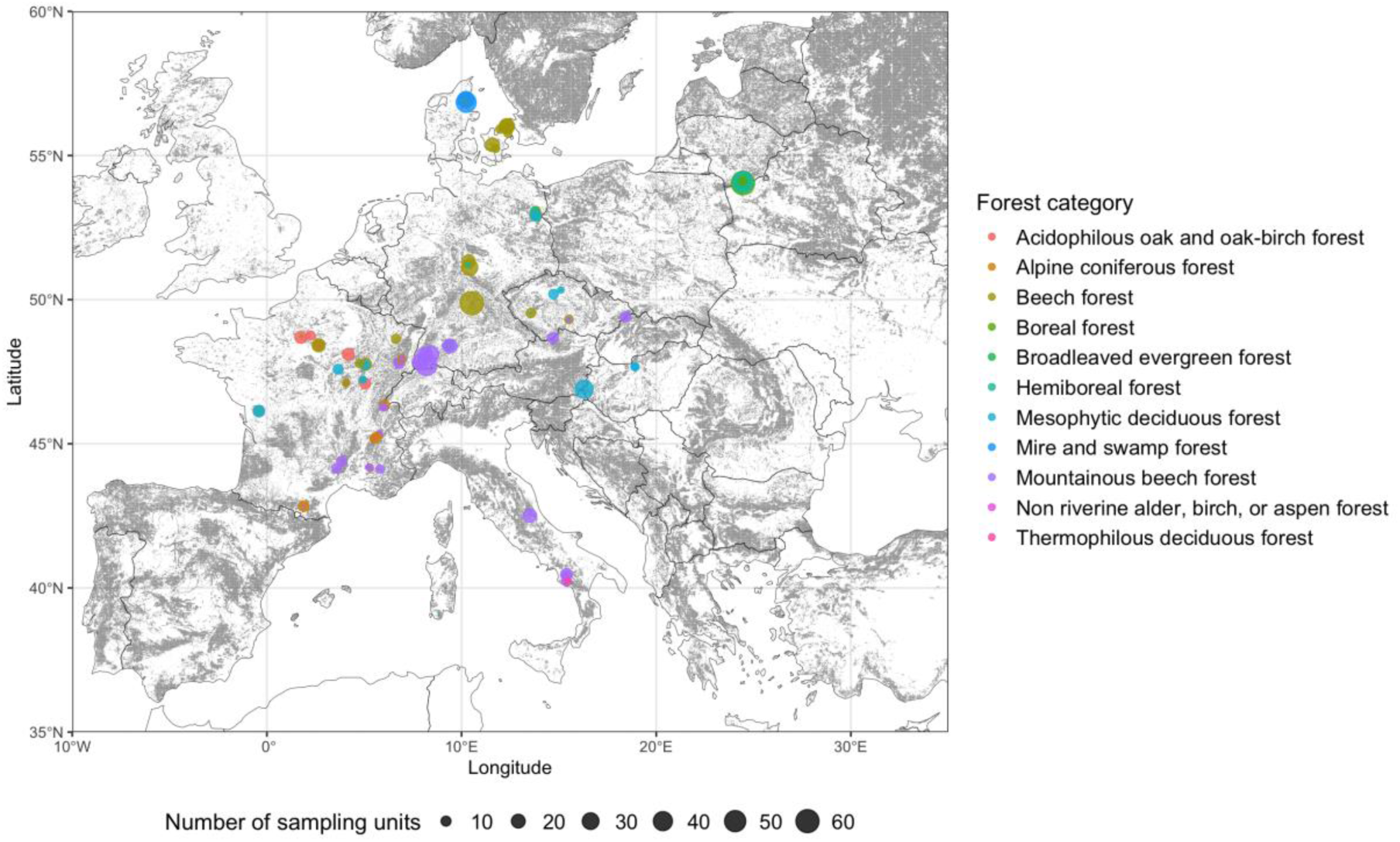
Spatial distribution of the sampling sites. Gray areas are covered by forests with a tree cover greater than 40% according to Kempeneers et al. (2011). The size of the dot is proportional to the number of sampling units. Colors correspond to forest categories (see Appendix 1 for details).

### 2.2. Indicators’ calculation

We used the State of Europe’s Forests (Forest Europe 2020) definitions to calculate the corresponding metrics for the indicators included in the Criterion 4 (Table 2). Since it was necessary to “translate” the definition into calculable values to analyze the relationship between indicator values and *indicanda* (i.e. scaled species richness), we proceeded as follows:

- indicator 4.6 *Genetic resources* was excluded since it was not possible to translate this indicator at the sampling unit or site levels;
- indicators 4.1 *Diversity of tree species*, 4.4 *Introduced tree species* and 4.5 *Deadwood* were directly calculated from field measurements. Instead of introduced tree species, we considered only invasive non-native woody species that can have a significant impact on forest biodiversity following Campagnaro et al. (2018), namely *Acer negundo*, *Ailanthus altissima, Prunus serotina*, *Quercus rubra* and *Robinia pseudoacacia*. We limited to these species since there was no regional reference list for introduced tree species and it would have been far beyond the scope of this study to create such a list, e.g. Norway spruce (*Picea abies*) is introduced in lowland France and Germany but native in mountains where it has also been planted in some places. In addition, no reference list of invasive tree species in Europe was available in the Forest Europe (2020) report;
- indicators 4.2 *Regeneration* and 4.3 *Naturalness* were derived from declarative metadata when merging the database (Burrascano et al. 2023). In the 4.3, forests were considered ‘undisturbed by man’ when declared without intervention by the data holder (i.e. ‘abandoned’, ‘unmanaged’, ‘primeval’). Time since the last intervention was not used here since it was not homogeneously declared. ‘Semi-natural’ refers to forests that were under any type of forest management regime (from even-aged to uneven-aged silvicultural systems), but with natural regeneration processes of trees. ‘Plantation’ forests overlapped with sites where regeneration derived from ‘Planting’ activities;
- indicators 4.8 *Threatened forest species* and 4.10 *Common forest bird species* were derived from biodiversity sampling and reference lists. For 4.8, we used the IUCN Red List (https://www.iucnredlist.org/) and calculated the total number of threatened species (categories VU, EN and CR) per sampling unit all groups together. For 4.10, we calculated the total number per sampling unit of the 34 forest birds classified by the Pan-European Common Birds Monitoring Scheme (https://pecbms.info/trends-and-indicators/indicators/);
- indicator 4.7 *Forest fragmentation* was calculated following the protocol applied by European Commission et al. (2019) on the Corine Land Cover 2018 layer (https://efi.int/knowledge/maps/forest). The forest area density (FAD) at fixed observation scale was obtained calculating the percentage of forest pixels in a 500 ha circular buffer, centered on each sampling unit;
- indicator 4.9 *Protected forests* was calculated based on the map of protected areas in Europe provided by the European Environment Agency (Nationally Designated areas
- CDDA: https://www.eea.europa.eu/data-and-maps/data/nationally-designated-areas-national-cdda-17). We extracted the proportion of Categories Ia, Ib and II in a 500ha circular buffer around each sampling unit. We then added these three values so that the total proportion may be higher than 100.

**Table 2:** Forest Europe’s biodiversity indicators definitions and data sources used to calculate them in this paper.

| Biodiversity indicators | Definition (Forest Europe 2020) | Data source | Metrics |
| --- | --- | --- | --- |
| 4.1. Diversity of tree species | Area of forest and other wooded land, classified by number of tree species occurring | Field measurements | Number of tree species per sampling unit |
| 4.2. Regeneration | Total forest area by stand origin and area of annual forest regeneration and expansion (natural regeneration, planting, coppicing) | Metadata | 3 classes:<br>Natural regeneration<br>Planting<br>Coppicing |
| 4.3. Naturalness | Area of forest and other wooded land by class of naturalness (undisturbed by man, semi-natural, plantations) | Metadata | 3 classes:<br>Unmanaged<br>Semi-natural<br>Plantation |
| 4.4. Introduced tree species | Area of forest and other wooded land dominated by introduced tree species<br>⇒ modified to invasive ligneous species (Campagnaro et al. 2018) | Field measurements | Number of invasive tree species per sampling unit |
| 4.5. Deadwood | Volume of standing deadwood and of lying deadwood on forest and other wooded land | Field measurements | Deadwood volume per ha |
| 4.6. Genetic resources | Area managed for conservation and utilization of forest tree genetic resources (in situ and ex situ genetic conservation) and area managed for seed production | Not assessed |  |
| 4.7. Forest fragmentation (FAD) | Area of continuous forest and of patches of forest separated by non-forest lands (European Commission et al. 2019) | European Forest Institute map of forest cover | Forest area density (FAD) within 500ha around the sampling unit |
| 4.8. Threatened forest species | Number of threatened forest species, classified according to IUCN Red List categories, in relation to total number of forest species | IUCN Red List species list | Species richness of threatened species (categories VU, EN and CR) |
| 4.9. Protected forests | Area of forest and other wooded land protected to conserve biodiversity, landscapes and specific natural elements, according to MCPFE categories | CDDA map of protected areas | Cumulative percentage of Ia, Ib and II categories areas |
| 4.10. Common forest bird species | Occurrence of common breeding bird species related to forest ecosystems | PECBMS | Species richness of forest birds |

### 2.3. Statistical analyses

We processed all analyses in R v.4.3.0 (R Core Team 2023). We used the scaled species richness of each taxonomic and functional group as a response variable. We fitted a generalized linear mixed model (GLMM) with site as a random effect on the intercept to account for the fact that two sampling units from the same site were more likely to be similar than two sampling units from different sites. We used the beta error distribution with logit link since standardized richness was continuous, strictly positive with a maximum value below one. We scaled continuous explanatory variables to improve the convergence and to make the estimates of the models comparable (“scale” function in R, which subtracts the standard deviation to the value and divides by the mean of the dataset). We used the glmmTMB function in the glmmTMB package (Brooks et al. 2017).

We processed the data in three steps:

- first, for each group, we fitted nine single models plus the null (intercept only) model with each indicator as an explanatory variable (Table 2). We compared the magnitudes of all single effects in the models and their significance and represented this using a heatmap of the standardized effects (package ggplot2, Wickham 2016). We tested differences between estimates of categorical variables (e.g. 4.3.naturalness) using Tukey post-hoc test. To search for potential non-linear relationships between indicators and *indicanda*, we also fitted generalized additive mixed models (GAMMs) with indicators as smoothers (package mgcv, Wood 2023). We found very few differences between GLMMs and GAMMs and preferred to stick to the former (comparisons are presented in Appendix 3).
- second, we assessed the relative performance of all single indicators by comparing the Akaike Information Criterion corrected for small samples (AICc, Burnham and Anderson 2002) of all single models including a null (intercept only) model.
- third, we searched for the best and most parsimonious linear combination (no interaction) of indicators that represented biodiversity based on AICc using the dredge function (package MuMIn, Barton 2023). In this process, we discarded the indicator 4.2. Regeneration since it was collinear with 4.3. Naturalness and a model containing both variables could not be fitted. When two competing models had a difference in AICc less than 2 points, we chose the most parsimonious one.

For all models, we calculated a pseudo-R^2^ (function r.squaredGLMM, package MuMIn) following Nakagawa et al. (2017) as a measure of the variance explained by a given model. Within a GLMM framework, pseudo-R^2^ comprises two components: a marginal R^2^ (R^2^m) which stands for the variance explained by the fixed effects only and a conditional R^2^ (R^2^c) which stands for the overall variance explained conditional to the random effects. Note that R^2^m of 0.2 to 0.4 are generally considered as an excellent fit (McFadden 1979).

## 3. Results

### 3.1. Data and indicators’ distribution

The most represented group in the database was tracheophytes (838 sampling units) followed by birds (825 sampling units, Table 3, see Burrascano et al. 2023 for a full description of the biodiversity dataset). All taxonomic and functional groups have been inventoried in at least 400 sampling units (Table 1). The distribution in the classes Regeneration (4.3.) and Naturalness (4.4.) were, however, strongly unbalanced (see also Table 1): the majority of sampling units were associated with “natural regeneration”, while “coppicing” - and “planting” to a lesser extent - were underrepresented; the majority of sampling units were within semi-natural forests, but the distribution was more balanced than for regeneration types. For quantitative indicators, the values taken were relatively balanced between groups (Table 3) and no strong collinearity was observed (see Appendix 4).

**Table 3:** Range of values for each indicator distributed over the different taxonomic and functional datasets (mean per sampling unit (sd) [min - max]). FAD = Forest Area Density.

| **Taxonomic and functional groups datasets** | **I4.1.tree.sp**<br>(species number) | **I4.4.invasive**<br>(species number) | **I4.5.deadwood**<br>(m<sup>3</sup>/ha) | **I4.7.fragmentation (FAD)**<br>(%) | **I4.8.threat.sp**<br>(species number) | **I4.9. Protected forests**<br>(%) | **I4.10.birds**<br>(species number) |
| --- | --- | --- | --- | --- | --- | --- | --- |
| Tracheophytes<br>(n = 838) | 1.96(2.06)<br>[0-11] | 0.01(0.08)<br>[0-1] | 41.83(60.72)<br>[0-444.89] | 0.85(0.16)<br>[0.13-1] | 0.25(0.56)<br>[0-3] | 14.92(33.66)<br>[0-142.4] | 3.63(1.78)<br>[1-11] |
| Bryophytes<br>(n = 422) | 1.91(1.84)<br>[0-11] | 0(0.06)<br>[0-1] | 39.15(55.86)<br>[0-461.41] | 0.84(0.18)<br>[0.06-1] | 0.2(0.5)<br>[0-3] | 27.02(43.16)<br>[0-142.4] | 4.41(2.63)<br>[1-15] |
| Beetles<br>(n = 509) | 1.96(2.01)<br>[0-11] | 0(0.07)<br>[0-1] | 41.1(57.2)<br>[0-444.89] | 0.83(0.17)<br>[0.08-1] | 0.26(0.59)<br>[0-5] | 16.63(35.16)<br>[0-142.4] | 4.15(2.43)<br>[1-12] |
| Birds<br>(n = 825) | 2.49(2.18)<br>[0-11] | 0.01(0.08)<br>[0-1] | 42.03(64.56)<br>[0-461.41] | 0.85(0.16)<br>[0.13-1] | 0.29(0.6)<br>[0-3] | 15.92(34.06)<br>[0-142.4] | 4.14(2.38)<br>[1-15] |
| Fungi<br>(n = 604) | 1.47(1.95)<br>[0-11] | 0.01(0.08)<br>[0-1] | 39.73(57.98)<br>[0-461.41] | 0.82(0.8)<br>[0.08-1] | 0.17(0.49)<br>[0-3] | 14.55(34.22)<br>[0-100] | 4.32(2.83)<br>[1-15] |
| Lichens<br>(n = 434) | 1.93(1.85)<br>[0-11] | 0(0.06)<br>[0-1] | 39.28(56.21)<br>[0-461.41] | 0.84(0.18)<br>[0.06-1] | 0.21(0.53)<br>[0-5] | 26.65(42.93)<br>[0-142.4] | 4.43(2.62)<br>[1-15] |

### 3.2. Links between single indicators and indicanda

Several of the Forest Europe indicators had significant relationships with the scaled species richness of one or more of the six taxonomic and functional groups (Figure 2, see Appendix 5 for values of the estimates).

**Figure 2:**
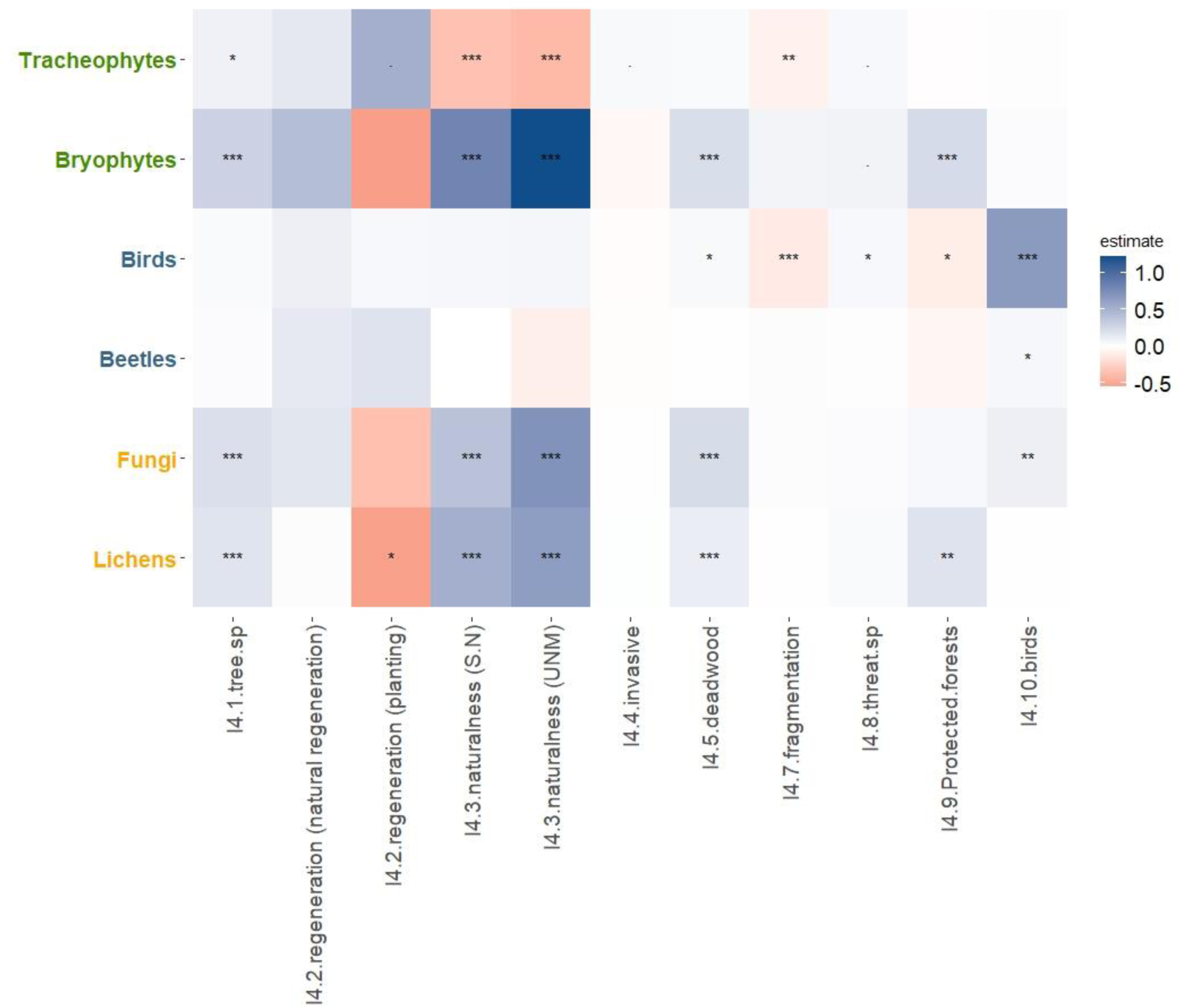
Heatmap representing the standardized estimates (slope) of scaled species richness obtained with generalized mixed models with Beta error distribution and logit link, using Forest Europe indicators as independent predictors. S.N = semi-natural, UNM = unmanaged. Intercept (reference) for 4.2 Regeneration is the “Coppice” class, and “Plantations” for 4.3 Naturalness (. p<0.1, * p<0.05, ** p<0.01, *** p<0.001).

Deadwood (4.5, Figure 3) and Diversity of tree species (4.1) showed four positive and (marginally) significant but generally weak relationships with tracheophytes, bryophytes, fungi and lichens (Appendix 5). Other indicators showed at least three significant (or marginally significant) relationships, with varying magnitudes, i.e. Fragmentation (FAD) (4.7), Protected areas (4.9, Figure 4) and Forest Birds (4.10). Surprisingly, the proportion of protected areas around sampling plots had a negative effect on birds and no effect on tracheophytes and fungi. Regarding Naturalness (4.3, Figure 5), unmanaged forest and semi-natural forests showed higher levels species richness in fungi, lichens and bryophytes compared to plantations. The opposite was true for tracheophytes. Threatened species (4.8) showed very few significant results (marginally positive for tracheophytes and bryophytes, positive for birds) while Invasive species (4.4) only showed a marginally significant positive relationship with tracheophytes. Plantation (4.2) showed a very strong negative relationship with lichens only, and a marginally significant negative effect on tracheophytes, while there were no differences between coppice and natural regeneration (Appendix 5). Marginal R-square values were generally low for single variables models (below 0.1) and most of the variations were captured by the random effect (conditional R-squares > 0.4, see Figures 3-5).

**Figure 3:**
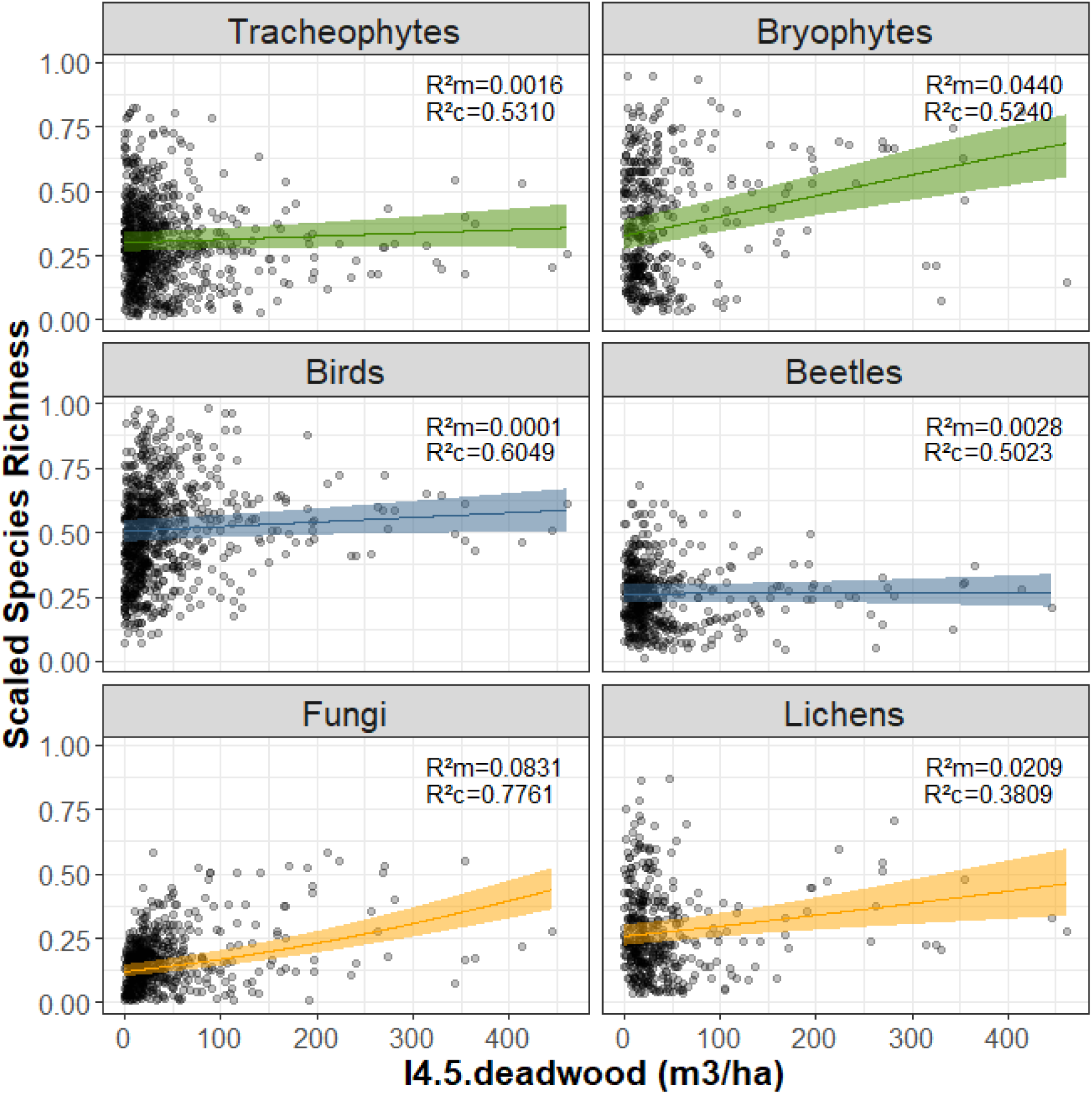
Relationships between scaled species richness of 6 taxonomic and functional groups and Forest Europe indicator 4.5. Total deadwood volume. Estimations are issued from generalized linear models with Beta error distribution and logit link. Plain line represents the mean estimate, ribbons the 95% confidence interval. R^2^m and R^2^c are marginal R-square for fixed effects, and conditional R-square with random effects, respectively.

**Figure 4:**
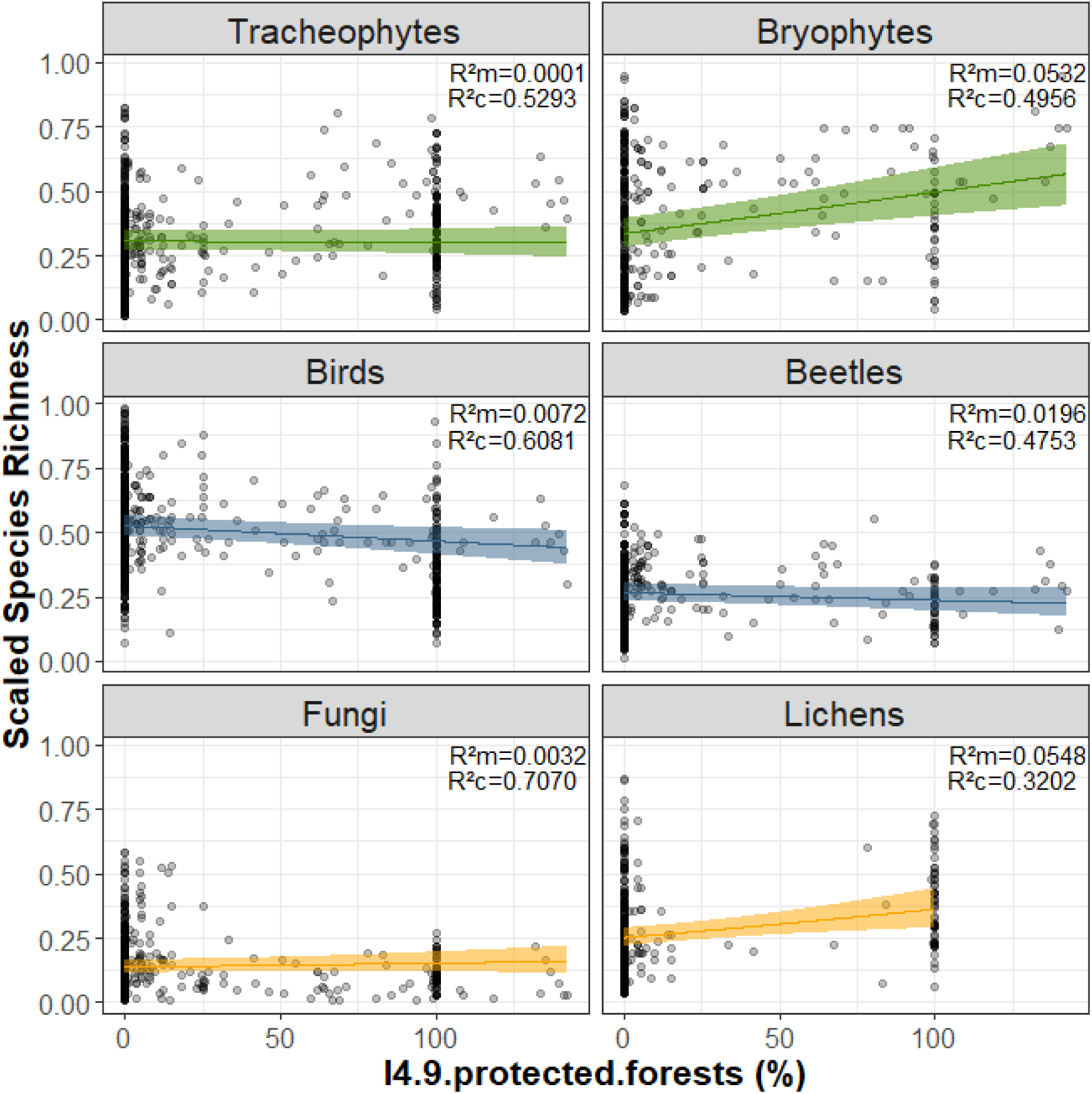
Relationships between standardized species richness of 6 taxonomic and functional groups and Forest Europe indicator 4.9 (Proportion of protected areas in a 500ha buffer). Estimations are issued from generalized linear models with Beta error distribution. Plain line represents the mean estimate, ribbons the 95% confidence interval. R^2^m and R^2^c are marginal R-square for fixed effects, and conditional R-square with random effects, respectively

**Figure 5:**
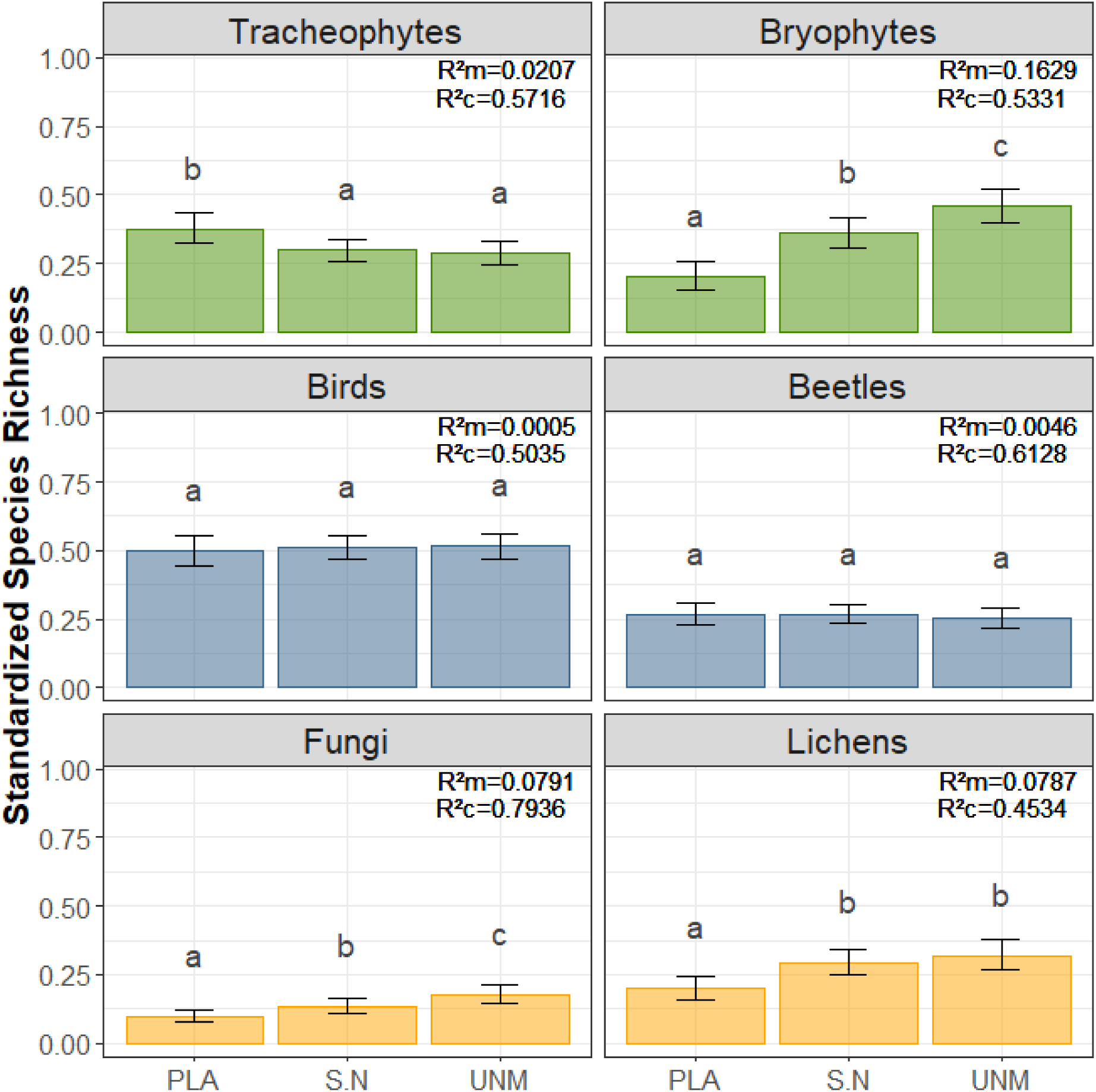
Relationships between standardized species richness of 6 taxonomic and functional groups and Forest Europe indicator 4.3. Naturalness: PLA = Plantations; S.N = Semi – Natural forests; UNM = Unmanaged forests. Estimations are issued from generalized linear models with Beta error distribution and logit link. Barplots represent mean estimates, error bars are the 95% confidence intervals. Letters indicate significant differences per group based on Tukey post-hoc test. R^2^m and R^2^c are marginal R-square for fixed effects, and conditional R-square with random effects, respectively

### 3.3. Relative performance of the different indicators

Comparing the AICc (Table 4) for all single models revealed that the indicator based on forest Naturalness (4.3) best explained the scaled species richness for three out of six groups (tracheophytes, bryophytes, and lichens); Regeneration (4.2) was within 2 points of AICc for tracheophytes and lichens. For birds and beetles, the indicator based on Forest birds (4.10) was the best explanatory one, while for fungi, Deadwood (4.5) stood first. Also note that the null model was never the best one and more than 2 AICc points away from the best.

**Table 4:**
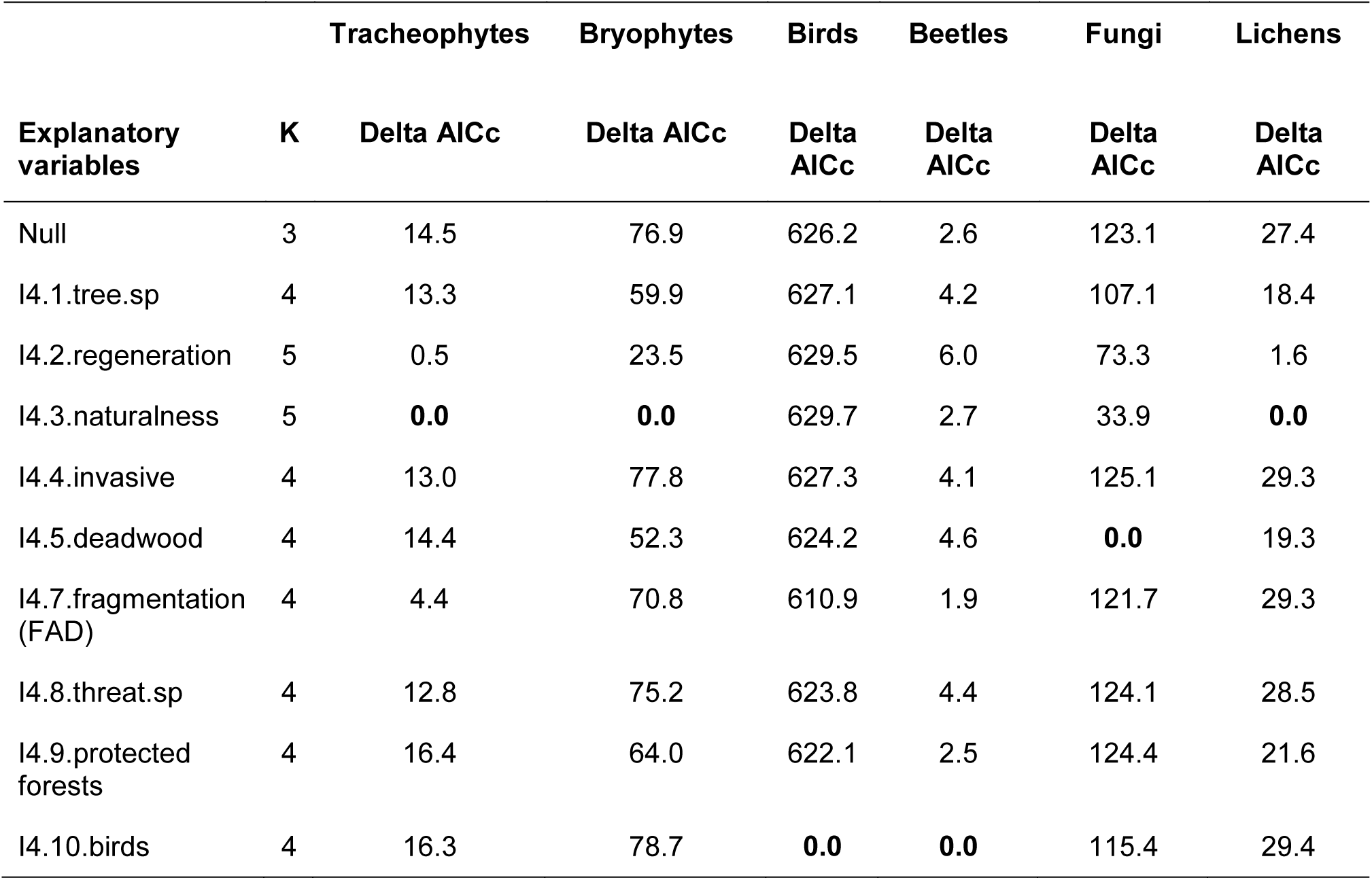
Differences of Akaike Information Criteria corrected for small samples (Delta AICc) for all generalized mixed models with Beta error distribution and logit link. K is the number of parameters in each model. The best model (delta AICc = 0) is in bold.

### 3.4. Combination of indicators

We used data dredging to find the best linear (i.e. without interaction) combination of indicators reflecting the variations in species richness of each group. Beetles were represented by only one indicator (Forest Birds, 4.10.), all the other ones had a combination of 3 to 4 indicators in their best models (Table 5). In terms of indicators, Invasive species (4.4.) was never selected. Conversely, and as observed before, Diversity of tree species (4.1.), Naturalness (4.3.), Deadwood (4.5.) were selected four times, followed by Forest Birds (4.10, 3 times). Finally, it is interesting to note that, except for Fragmentation (FAD, 4.7.), all indicators had positive effects, and with the exception of beetles, all multivariate models performed better than univariate models containing only one indicator (in terms of AICc, they had at least 2 points less than single variable ones, Table 5).

**Table 5:** Scaled estimates of the models selected by data dredging combining all indicators from Forest Europe (without interaction). Estimates are issued from generalized linear mixed models with Beta error distribution and logit link. “+” indicates the presence of the factor in the model. The best model was the most parsimonious (lowest degrees of freedom - df) with the lowest Akaike Information Criterion corrected for small samples (AICc). “delta” indicates the AICc difference with the best model. logLik = logarithm of the likelihood. R^2^m and R^2^c are marginal R-square for fixed effects, and conditionnal R-square with random effects, respectively.

| **Taxonomic and functional groups** | **Tracheophytes** | **Bryophytes** | **Birds** | **Beetles** | **Fungi** | **Lichens** |
| --- | --- | --- | --- | --- | --- | --- |
| Intercept | -0.47 | -1.28 | 0.11 | -1.03 | -2.14 | -1.34 |
| I4.1.tree.sp | 0.11 | 0.17 |  |  | 0.12 | 0.12 |
| I4.3.naturalness | + | + |  |  | + | + |
| I4.4.invasive |  |  |  |  |  |  |
| I4.5.deadwood | 0.06 | 0.09 |  |  | 0.17 | 0.07 |
| I4.7.fragmentation (FAD) | -0.072 |  | -0.07 |  |  |  |
| I4.8.threat.sp |  |  | 0.05 |  |  |  |
| I4.9.protected forests |  |  |  |  |  |  |
| I4.10.birds |  |  | 0.65 | 0.05 | 0.08 |  |
| Df | 8 | 7 | 6 | 4 | 8 | 7 |
| AICc | -1246.18 | -384.61 | -1678.81 | -1123.87 | -1743.14 | -547.79 |
| Delta AICc | 0.61 | 1.42 | 0.19 | 0.96 | 1.23 | 1.69 |
| R2m | 0.071 | 0.341 | 0.650 | 0.009 | 0.132 | 0.150 |
| R2c | 0.828 | 0.856 | 1.000 | 0.808 | 0.847 | 0.697 |
| AICc (best single variable models) | -1226.72 | -375.82 | -1648.93 | -1123.87 | -1671.48 | -541.632 |

Marginal R-squares were higher than those observed for univariate models, reaching 0.34 for bryophytes (Table 5). Conditional R-squares were even stronger than those observed for univariate models (all >0.8 except for lichens, Table 5).

## 4. Discussion

We analyzed the links between Forest Europe’s biodiversity indicators and the scaled richness of six taxonomic and functional groups on a unique multi taxonomic dataset covering most temperate forest types in Europe (Burrascano et al. 2023, Appendix 1). We showed that these indicators perform unequally: while some correctly predicted the biodiversity of several groups, no one acted as a universal indicator, questioning their strength to predict biodiversity in European forests. In contrast, our results reinforced the approach based on multiple complementary indicators of the same *indicandum*. This also raises the question of contextualization of these indicators, to better assess state and trends of biodiversity across Europe, and opens perspectives for improvement.

### 4.1. Performance of current forest biodiversity indicators

While evidence on the link between some indicators and the biodiversity they are supposed to indicate remains incomplete (Gao et al. 2015; Penone et al. 2019; Zeller et al. 2023), our results highlighted several habitat-species relationships and habitat preferences of different groups at different scales.

#### 4.1.1. Local scale indicators (sampling unit)

Deadwood volume, followed by diversity of tree species, were two indicators that often correlated with scaled species richness, both in univariate and multivariate analyses. Deadwood and diversity of tree species have complementary roles for biodiversity (Storch et al. 2023). Deadwood provides a substrate and a resource for numerous species that depend on it for part of their life cycle (Lassauce et al. 2011; Müller et al. 2019; Müller et al. 2015; Parajuli and Markwith 2023), but also for facultative species (Graf et al. 2022). Indeed, the richness of bryophytes, fungi and lichens, and to a lesser extent birds, correlated positively to deadwood volume in our analyses. Surprisingly however, saproxylic beetles richness did not correlate significantly to deadwood volume despite a weak positive effect. It is likely that, for this group in particular, deadwood does not act as a universal indicator that could be transposed in all situations (see also Zeller et al. 2023).

Diversity of tree species also showed positive correlation with the biodiversity of several taxonomic and functional groups (bryophytes, fungi, lichens and more marginally tracheophytes). Different tree species provide different habitat conditions for epiphytic and saproxylic species living in forests (e.g. Boch et al. 2021; Cavard et al. 2011; Leidinger et al. 2021). These conditions encompass direct biotic interactions, different chemical properties of the bark and wood, decomposition rates as well as differential light interception (Gosselin et al. 2017; Zeller et al. 2023). This in turn provides niche heterogeneity for several species and thus enhances biodiversity and confirms the role of diversity of tree species as a biodiversity indicator (see however Likulunga et al. 2021; Wildermuth et al. 2024).

Invasive non-native tree species did not show any correlation with biodiversity, apart from a marginal positive effect on tracheophytes supporting high local tracheophyte diversity to be often an indicator of disturbance. Despite the negative effects of invasive non-native species introduction on native *flora*, *fauna* and *mycota*, this effect is rather limited in our dataset since it contains very few invasive species in total (maximum one tree species per plot) reflecting that no projects targeted the effects of plantations rich in, or dominated by, invasive non-native tree species on biodiversity. This limited gradient probably does not allow to show significant effects on biodiversity. In addition, an indicator on introduced species rather than invasive ones – as defined in Forest Europe’s indicators – may perform better.

#### 4.1.2. Stand scale indicators (management unit)

Regeneration origin (4.2) and Naturalness (4.3) had a strong effect on the biodiversity of the six groups. The biodiversity of bryophytes and fungi was higher in unmanaged forests 24 compared to semi-natural forests, and, including lichens, higher than in plantation forests (Figure 5). On the contrary, plantations had a marginal positive effect on tracheophytes compared to semi-natural and unmanaged forests, and naturalness had no effects on birds and beetles. The negative effects of plantations and the positive effects of management abandonment or primeval forests have been thoroughly documented (e.g. Chaudhary et al. 2016; Langridge et al. 2023; Paillet et al. 2010). However, our results should be nuanced by the fact that the distribution of the data in the different categories of regeneration and naturalness were strongly unbalanced, with only few plots located in coppice-managed forests and a vast majority in semi-natural forests (Table 1). That said, unmanaged forests had a generally positive influence on biodiversity in our dataset, especially for deadwood dependent species (epixylic bryophytes and lichens but not for saproxylic beetles), while plantations had detrimental effects on several other groups. Indeed, habitat provision and continuity are higher in semi-natural forests, not to speak of unmanaged forests, which allow the persistence of dispersal limited species (e.g. Boch et al. 2013; Boch et al. 2021). The positive response of tracheophytes in plantations may seem surprising, but probably reflects a higher share of disturbance tolerant herbs in more disturbed sites, as shown in several individual studies (Atrena et al. 2024; Boch et al. 2013). In addition, the semi-natural category encompasses a wide range of management types (Trentanovi et al. 2023), and unmanaged forests in our dataset have mostly been recently abandoned, which may cause a decrease in the biodiversity of tracheophytes during the first decades after abandonment (e.g. Paillet et al. 2010).

#### 4.1.3. Landscape scale indicators (planning unit)

The proportion of protected forests around the plots positively influenced the richness of bryophytes and lichens, highlighting the positive effects of protection for these groups, but surprisingly had a negative effect on birds (see Honkanen et al. 2010). This may be related to the consideration of only strictly protected areas in this indicator. Strictly protected areas have no or little human interference, reducing possible heterogeneity at the landscape scale. Forest fragmentation showed negative effects on tracheophytes and birds, a positive effect on bryophytes, and marginally on beetles and fungi. It should be noted that since it is based on forest area density (European Commission et al. 2019), it represents forest cover rather than its discontinuity and configuration. Besides, the landscapes surrounding our sampling units presented a high forest cover (77% on average, in a 500 ha buffer, Table 1). Thus, the observed relationships reflect that forest area density could have been beneficial for forest interior species that are sensitive to edge effects (bryophytes or lichens), but detrimental to non-forest, open habitat or light demanding species, such as tracheophytes and open-habitat birds. Recent multitaxa studies proved the disproportionately high value of small patches, harboring richer assemblages, even when focusing only on protected species (Riva and Fahrig 2022). Besides, it has been highlighted that biodiversity conservation in human-modified forest landscapes is better achieved by maintaining at least 40% of forest cover, rather than focusing on fragmentation and configuration (Arroyo-Rodríguez et al. 2020). Hence, the role of fragmentation, as it is currently estimated, should probably be reconsidered to take into account both the amount of forest cover and the value of the patches.

#### 4.1.4. Indicators based on species data

Contrary to the previous indicators, threatened species and birds are based on direct biodiversity sampling (other than trees). These indicators directly describe the evolution of a small part of the biodiversity, but generally poorly indicate the richness of other taxa. Most of the results observed are linked to the fact that these indicators represent subsets of larger groups (evidently forest birds vs. birds). However, despite several pieces of evidence showing that congruence between taxa is generally small, especially in forests (Burrascano et al. 2018; Westgate et al. 2017), forest birds positively correlated to beetles and fungi. Probably these groups respond to the same favorable habitat conditions, but this was not reflected in the multivariate analyses.

### 4.2. Improving Forest Europe indicators to predict biodiversity patterns

#### 4.2.1. Combining indicators

Our study is one of the first to test Forest Europe indicators against multitaxonomic data at the European level. During the model selection process, the best univariate models were more often involving naturalness (tracheophytes, bryophytes and lichens) followed by forest birds (birds and beetles) and deadwood (fungi). However, the multivariate models did perform better than models including only single variable (except for beetles), and often the best ones combined several indicators to reflect variations of the *indicandum* (as shown by lower AICc and higher pseudo-R-squares values). Only beetles were best indicated by forest birds, but with a low magnitude, and birds by three combined indicators (including forest birds and threatened species). The magnitude of the effects individual values of the estimates in the multivariate models were comparable to those of single variable models (Appendix 5), which confirmed that indicators were not collinear and reflect different aspects of the biology of the target taxonomic and functional groups.

#### 4.2.2. Considering context and metrics

We limited our approach to the strict definition of the indicators as used in Forest Europe, but higher performance could probably be reached by at least two improvements. First, adding context to the indicators could probably reveal that they need to be adapted locally (Chiarucci et al. 2012; Honkanen et al. 2010). This is confirmed by the large variance absorbed by the random site effects in our models (Table 5). Context-related variables to be considered include elevation (mountain vs. lowlands), biome (Mediterranean, temperate, boreal) or European forest types.

Second, most of the metrics we used were abundance metrics (apart from tree, threatened and bird species indicators) that quantify habitat available for species. Indicators based on diversity of resources (following the heterogeneity-diversity theory, Tews et al. 2004) could perform better, e.g. in the case of deadwood and saproxylic beetles (Bouget et al. 2013), or tree-related microhabitats and birds and bats (Paillet et al. 2018). It would then be interesting to assess the performance of other indicator metrics vs. the current ones in assessing forest biodiversity.

On the biodiversity – *indicandum* – side, we studied only total scaled richness as a response variable, and evidently, the results may be different for other, more specialized groups, or other metrics of biodiversity (abundance, occurrence of individual species, functional diversity, e.g. Chianucci et al.). Such an approach remains to be tested (see e.g. Lelli et al. 2019) but was beyond the scope of the present study. In addition, some of the references we used were probably incomplete regarding some groups: e.g. almost no lichens are included in the list we used for red-listed species, but the proportion of threatened species at the national levels may be high.

#### 4.2.3. Future applications

Stevenson et al. (2021) claimed that indicators were often implemented without clear considerations of their purposes and utility in terms of decision-making. We argue that, while combinations of current Forest Europe indicators are useful to delineate general trends in biodiversity, they poorly reflect the patterns of species diversity. Adding context and analyzing the performance of other – more diversity-driven – metrics would help better reporting on biodiversity (e.g. Alterio et al. 2023; Paillet et al. 2018), in particular for groups for which the tested indicators showed a poor or a lack of predictive power (e.g., beetles).

Such improvements would also be beneficial to the use of indicators beyond general trends, to evaluate management and policy actions, decisions, or set biodiversity targets (Stevenson et al. 2021), also beyond forest ecosystems. For example, we showed that larger negative effects on biodiversity were observed in planted forests. This poses key challenges for making the 3 billion trees planting promoted by the European Forest Strategy for 2030 a beneficial action for biodiversity in forests (Sills et al. 2020), if high growing rate plantations such as introduced Douglas fir (*Pseudotsuga menziesii*) or *Eucalyptus* spp. are promoted against semi-natural forests. Conversely, promoting old-growth and unmanaged forests, as well as restoration of monocultures and conversions towards semi-natural forests, could have a positive effect on biodiversity. It is crucial to assess and balance these potential effects with the use of current data available and biodiversity indicators before taking actions or to modulate actions in favor of biodiversity against detrimental ones.

## 5. Conclusions: towards new indicators definitions and better reporting

Even if novel agreements, such as the Kunming-Montreal Global Biodiversity Framework (Convention on Biological Diversity 2022), continue stressing the importance of sustainable management in forestry, there is the risk that environmental policies could be biased towards market-related values (Pascual et al. 2023). Thus, the availability and application of reliable biodiversity indicators is both urgent and imperative.

Many forest biodiversity indicators are proxies based on pre-existing data mostly issued from National Forest Inventories (Tomppo et al. 2010) and more generally describe habitat attributes or area-based metrics rather than biodiversity *per se* (Marshall et al. 2020). Despite recent progresses based on international initiatives (namely the Essential Biodiversity Variables, GEOBON, IPBES), monitoring the state and trends of forest biodiversity solely based on proxies is not satisfactory: while proxies are generally easier to measure than species themselves, they are prone to demographic effects such as extinction debts or colonization credits. In other words, the presence of a given habitat – such as deadwood – does not guarantee the presence of the species that depend on it (e.g. Paillet et al. 2018). This results in a time lag between the effects of management or policy on actual biodiversity that may impair adaptive capacity in case of negative effects. In addition, the response of biodiversity to a given indicator depends on the taxonomic or functional group studied (Zeller et al. 2022), so no indicator may represent biodiversity overall, as it was confirmed by our analyses.

Even if proxies are useful and relatively easy to collect, they are probably not reactive enough to assess the state and dynamics of biodiversity. We thus recommend assessing both habitat features as well as species to clearly evaluate the effects of various pressures on biodiversity and being able to cope with them in an adaptive way.

## Acknowledgements

This work was funded by the EU Framework Programme Horizon 2020 through the COST Association (www.cost.eu): COST Action CA18207: BOTTOMS-UP – Biodiversity Of Temperate forest Taxa Orienting Management Sustainability by Unifying Perspectives. The authors are thankful to all those experts contributing to the data here harmonized and resumed that were not listed as data contributors.

## Appendices

## Appendix 1

Distribution of sampling units used for the analyses according to forest type (European Environment Agency 2007, Table 1.1), mean annual temperature (Figure 1.1) and mean annual precipitation (Figure 1.2) derived from Worldclim 2.0 (Fick and Hijmans 2017).

**Table 1.1:** Distribution of the studied sampling units in the 14 forest categories of the (European Environment Agency 2007)

| Forest category | Name | Tracheo<br>phytes | Bryophyt<br>es | Birds | Beetles | Fungi | Lichens |
| --- | --- | --- | --- | --- | --- | --- | --- |
| 1 | Boreal forest | 64 | 0 | 64 | 0 | 0 | 0 |
| 2 | Hemiboreal forest and nemoral coniferous and mixed broadleaved coniferous forest | 100 | 18 | 101 | 36 | 36 | 35 |
| 3 | Alpine coniferous forest | 25 | 18 | 25 | 21 | 16 | 0 |
| 4 | Acidophilous oak and oak-birch forest | 60 | 56 | 60 | 57 | 59 | 0 |
| 5 | Mesophytic deciduous forest | 83 | 72 | 83 | 68 | 69 | 47 |
| 6 | Beech forest | 194 | 103 | 180 | 171 | 191 | 128 |
| 7 | Mountainous beech forest | 253 | 105 | 254 | 107 | 176 | 166 |
| 8 | Thermophilous deciduous forest | 9 | 9 | 9 | 0 | 7 | 9 |
| 9 | Broadleaved evergreen forest | 1 | 0 | 0 | 0 | 1 | 0 |
| 14 | Floodplain forest | 49 | 41 | 49 | 49 | 49 | 49 |

**Figure 1.1:**
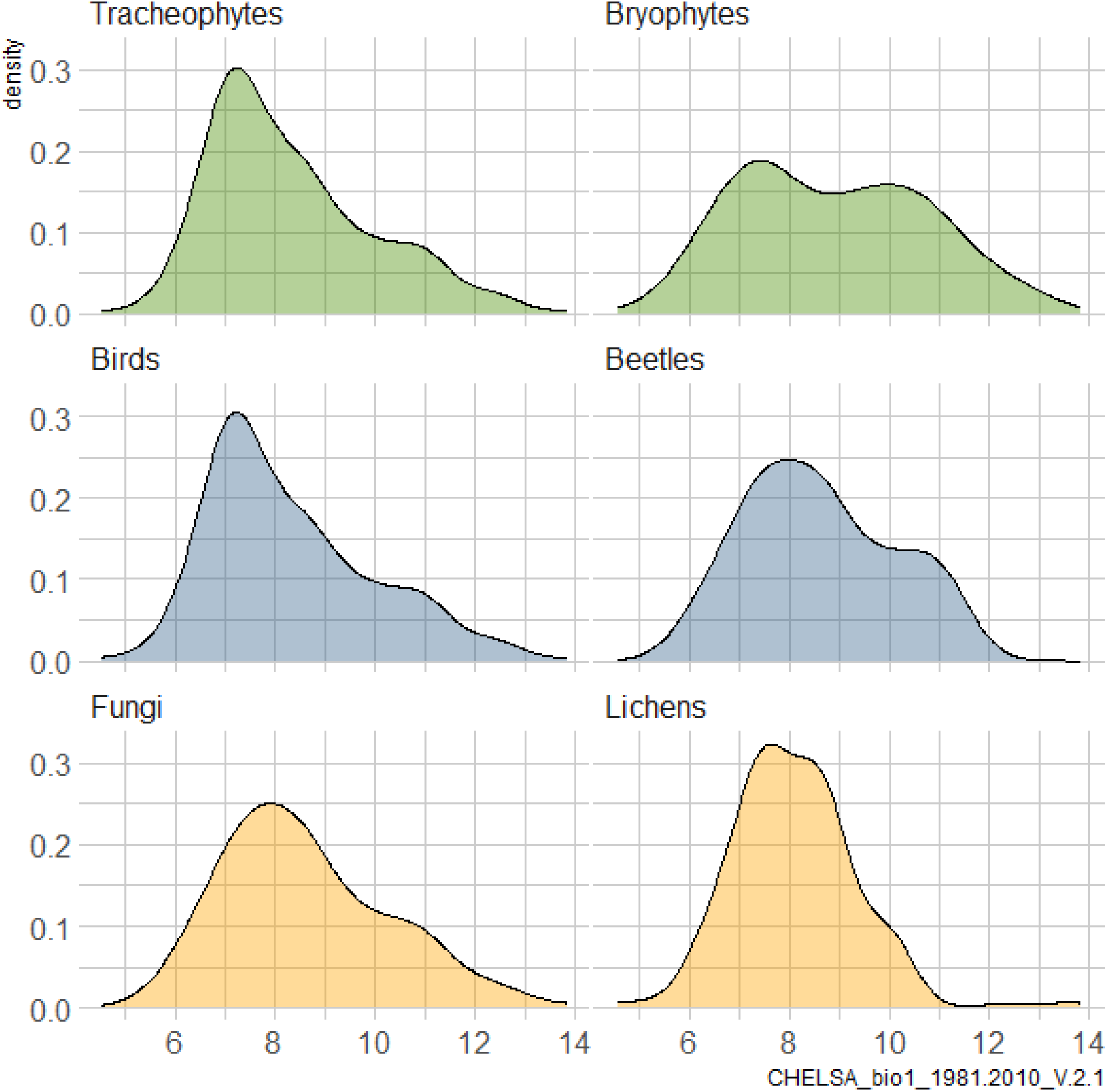
Mean annual temperatures for the different taxa datasets derived from Worldclim 2.1 (1991-2010).

**Figure 1.2:**
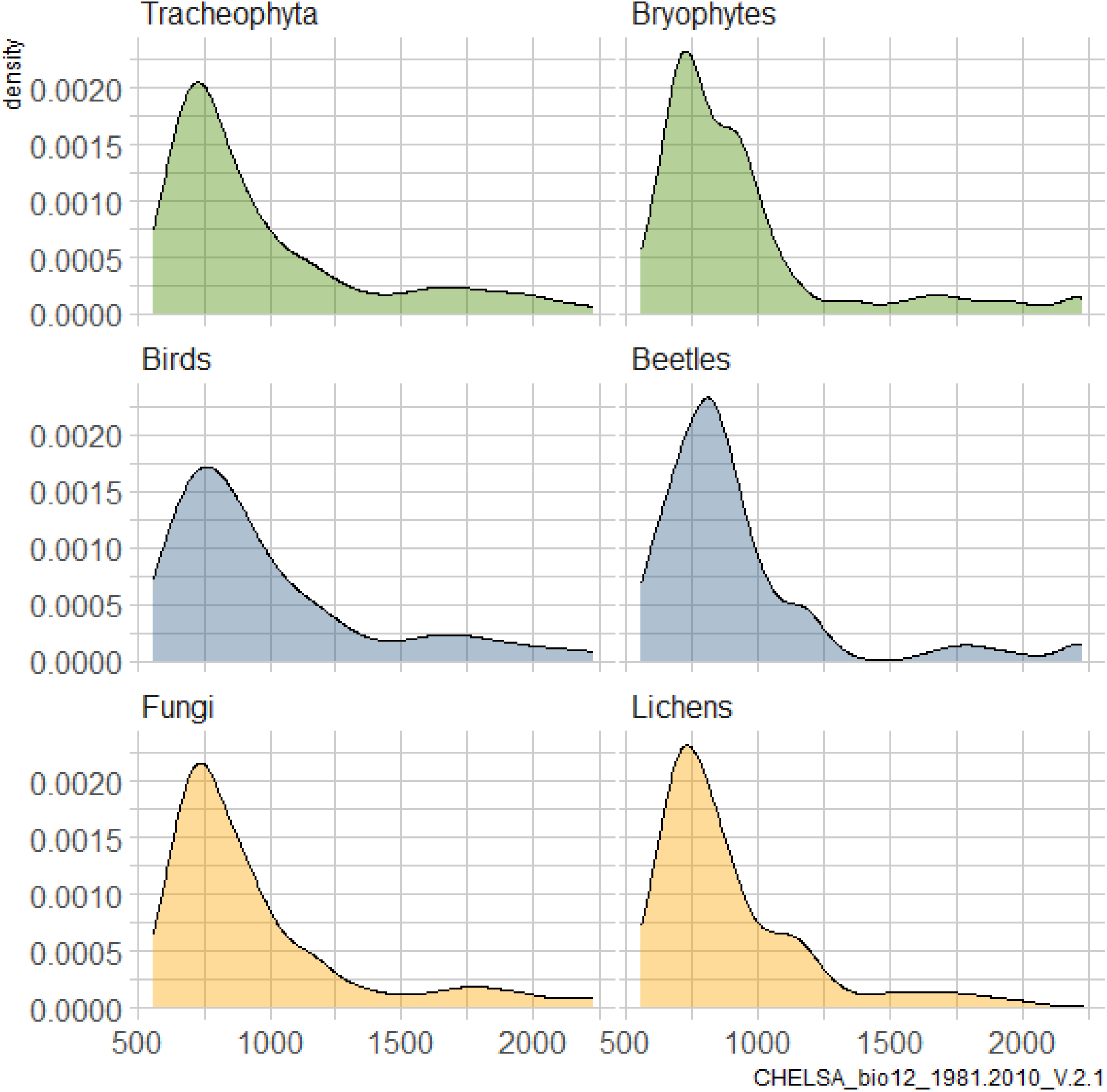
Mean annual precipitation for the different taxa datasets derived from Worldclim 2.1 (1991-2010)

## Appendix 2

Sensitivity analysis of the biodiversity response variable.

Analysing original biodiversity data from different sites (here 23 and 43 sites depending on the taxa) compiled from different studies (Burrascano et al. 2023) is challenging, as sampling intensity between studies varies and sites may have different regional biodiversity pools. For example, boreal forests usually show a lower tracheophyte richness than forests of warmer temperate climates. This may bias findings in both ways, i.e. falsely not detecting significant relationships and falsely claiming significant relationships.

In statistical regression analyses, mixed models with site as a random effect on the intercept can partly compensate for that kind heterogeneous data. However, mixed models only account for the effect of sampling intensity and regional biodiversity on the intercept and not the slope. This is why we standardized biodiversity, i.e. species richness, using three methods:

- Gamma standardization = alpha_Plot_ / gamma.asy_Site_ - (gamma.asy_Site_ is the Chao2 estimator)
- Alpha standardization = alpha_Plot_ / max(alpha_Plots in Site_)
- Z score standardization = (alpha_Plot_ - mean(alpha_Plots in Site_)) / SD(alpha_Plots in Site_)

Gamma standardization, the plot level species richness scaled to the asymptotic gamma richness at the site level, is used in the main text. Values are confined to the range 0 to 1, but because plot level biodiversity is lower than asymptotic gamma biodiversity values close to 1 were rarely observed. Also, species rich plots may be attributed with low values when located in species rich sites (Figure 2-1a).

Alpha standardization, the plot level species richness scaled to the maximum plot level species richness within a site, shows a more even distribution over the range of 0 to 1 than gamma standardization with one maximum biodiversity plot per site, but does not distinguish between species poor and species rich sites (Figure 2-1b).

The Z score standardization, which is widely used in ecology to compare effect sizes, resembles the method we used to scale our metric explanatory variables. This approach resulted in differing ranges for taxa (Figure ?-1c).

We fitted a generalized linear mixed model (GLMM) with site as a random effect to account for the spatial nesting of the data. We used beta error distribution with logit link for gamma and alpha standardized biodiversity, Gaussian error distribution for Z score biodiversity, and Poisson error distribution for raw biodiversity.

We found that the response pattern of the taxa to the explanatory variables was generally only slightly affected by the biodiversity metric employed (Figure ?-2). Results for gamma standardization, alpha standardization and raw biodiversity largely overlapped in response direction, response strength and significance levels. Z score standardization deviated from the other standardizations as a lower number of significant relationships, also showing a lower response strength, was found.

**Figure 2.1.**
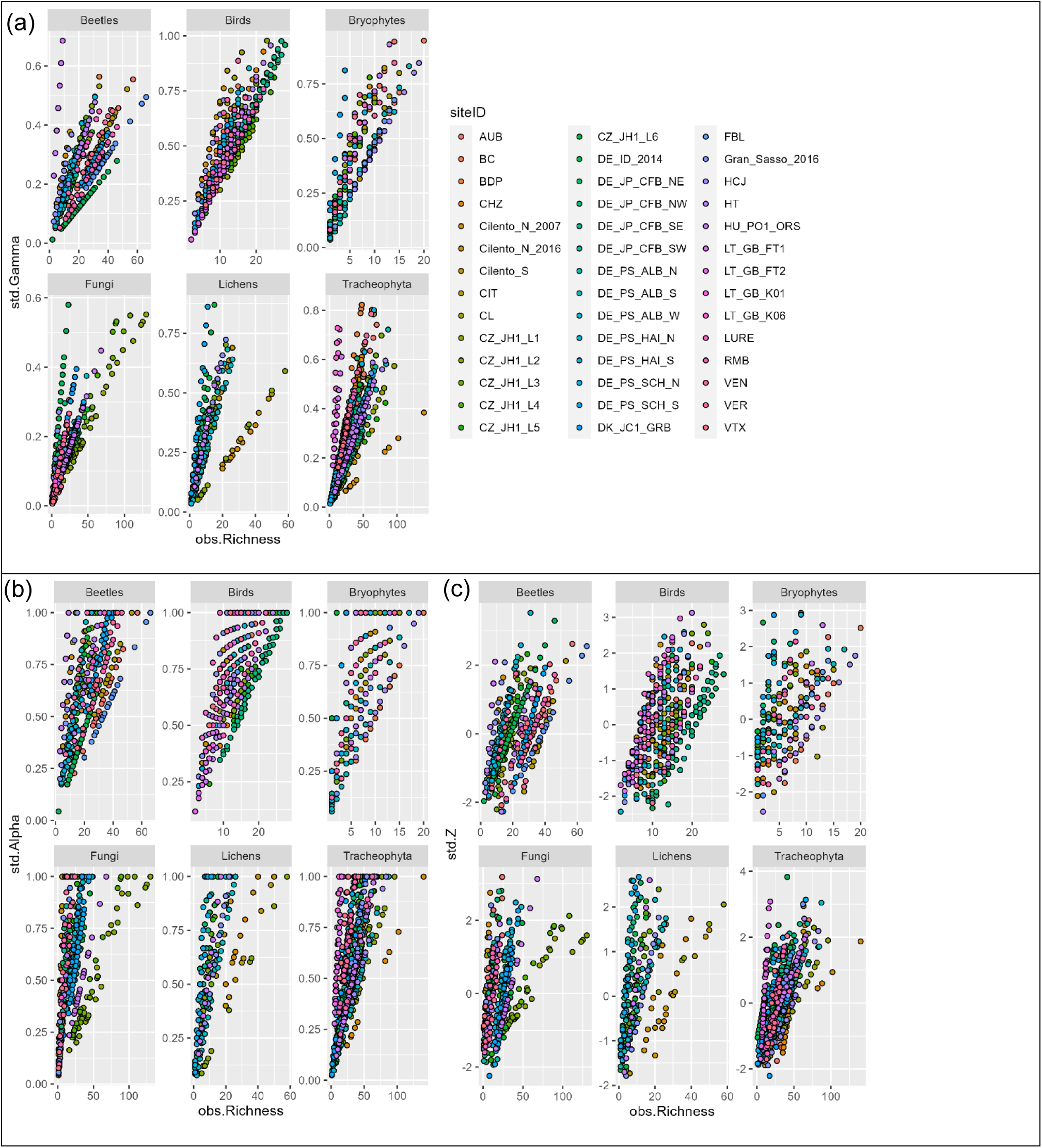
Scatter plot of raw biodiversity (species richness) versus gamma-standardized, alpha-standardized, Z-standardized biodiversity for sites.

**Figure 2.2.**
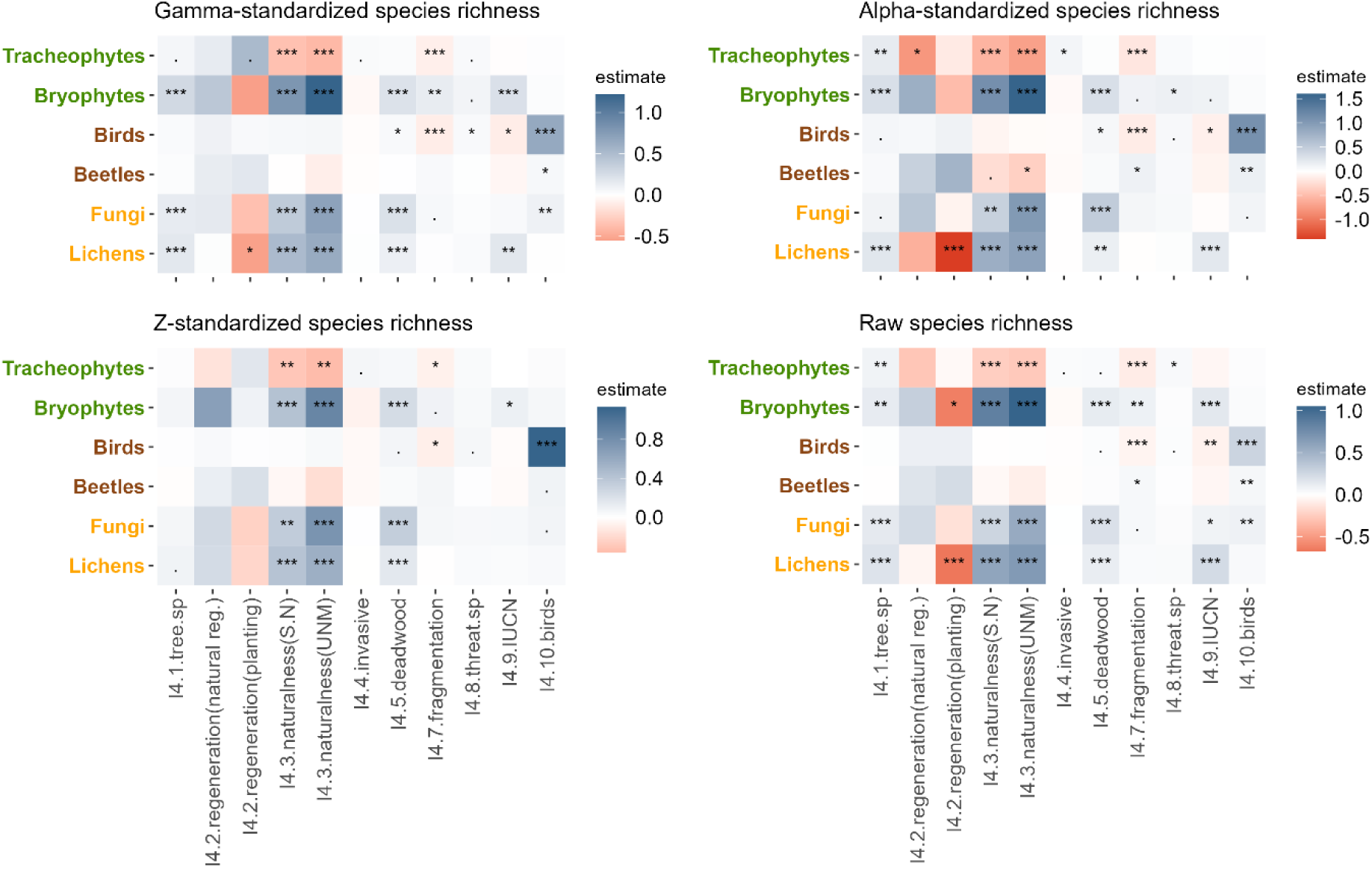
Overview of standardized estimates (slope) for four different biodiversity metrics. Results for gamma-standardized, alpha-standardized, Z-standardized and raw species richness obtained with generalized mixed models using Forest Europe indicators as independent predictors. S.N = semi-natural, UNM = unmanaged. Intercept (reference) for 4.2 Regeneration is the “Coppice” class, and “Plantations” for 4.3 Naturalness (p<0.1, * p<0.05, ** p<0.01, *** p<0.001).

## Appendix 3

Comparison of linear (GLMMs) vs non-linear (GAMMs) models analyses between standardized species richness and several Forest Europe’s biodiversity indicators.

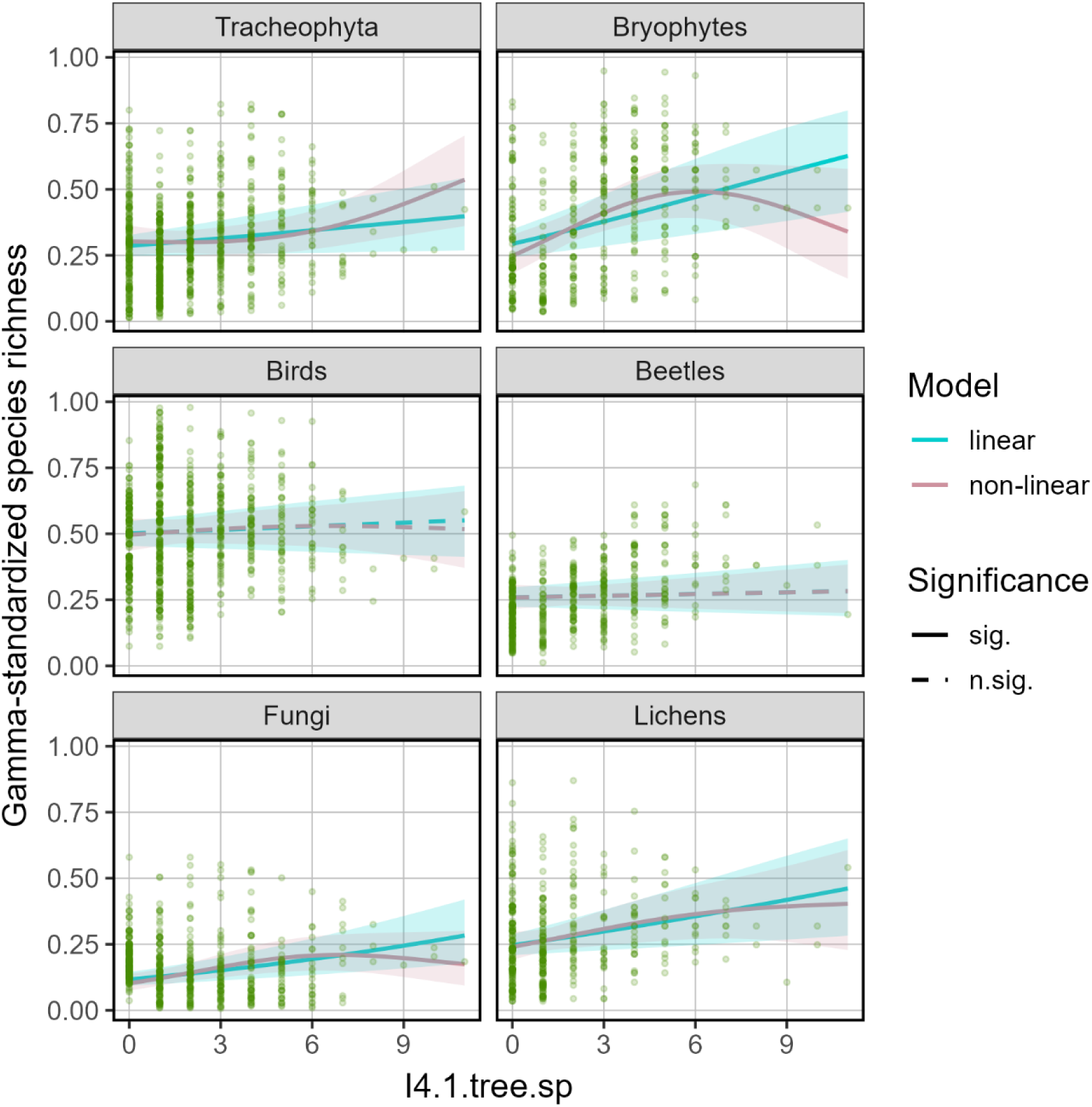

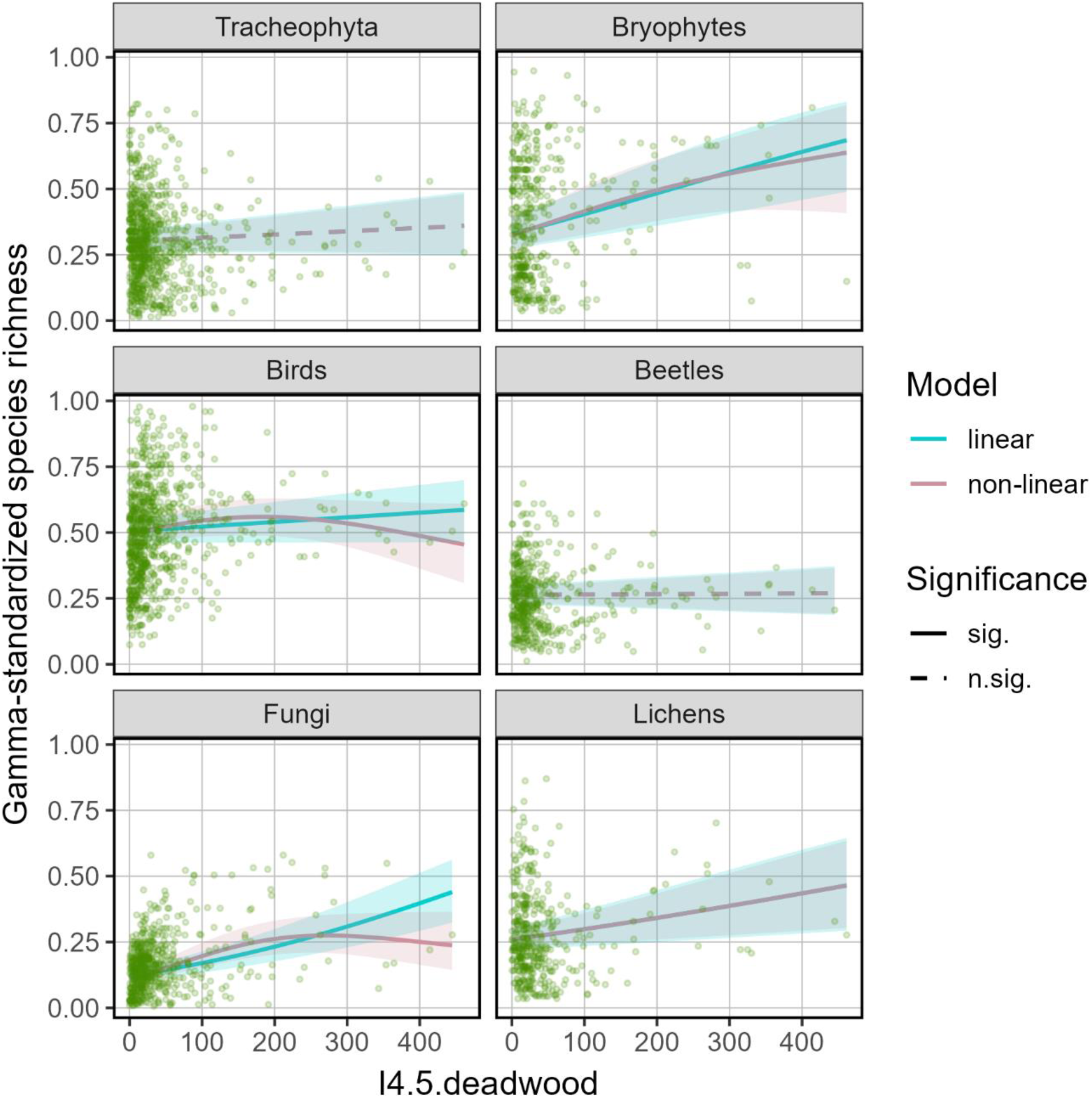

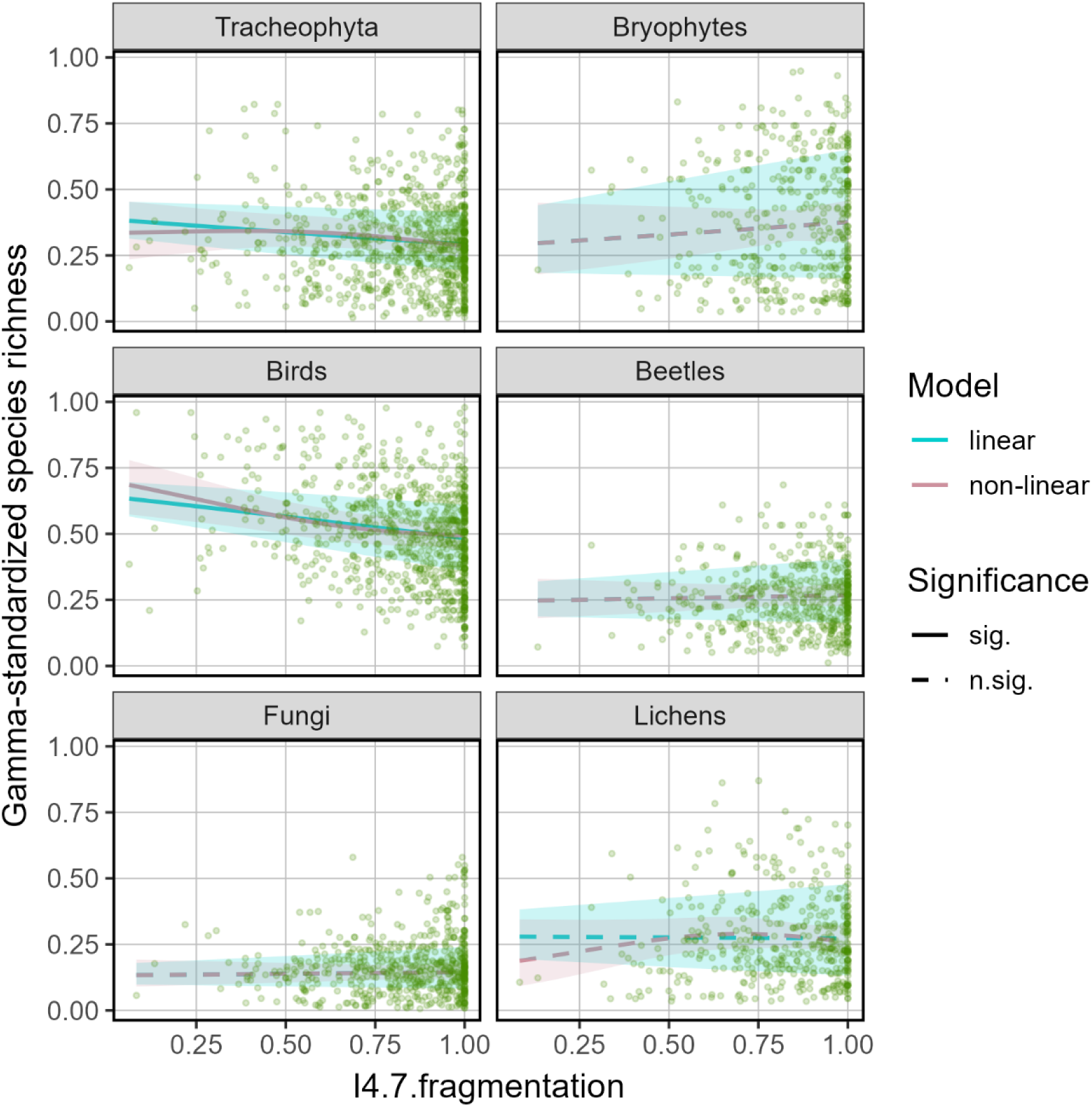

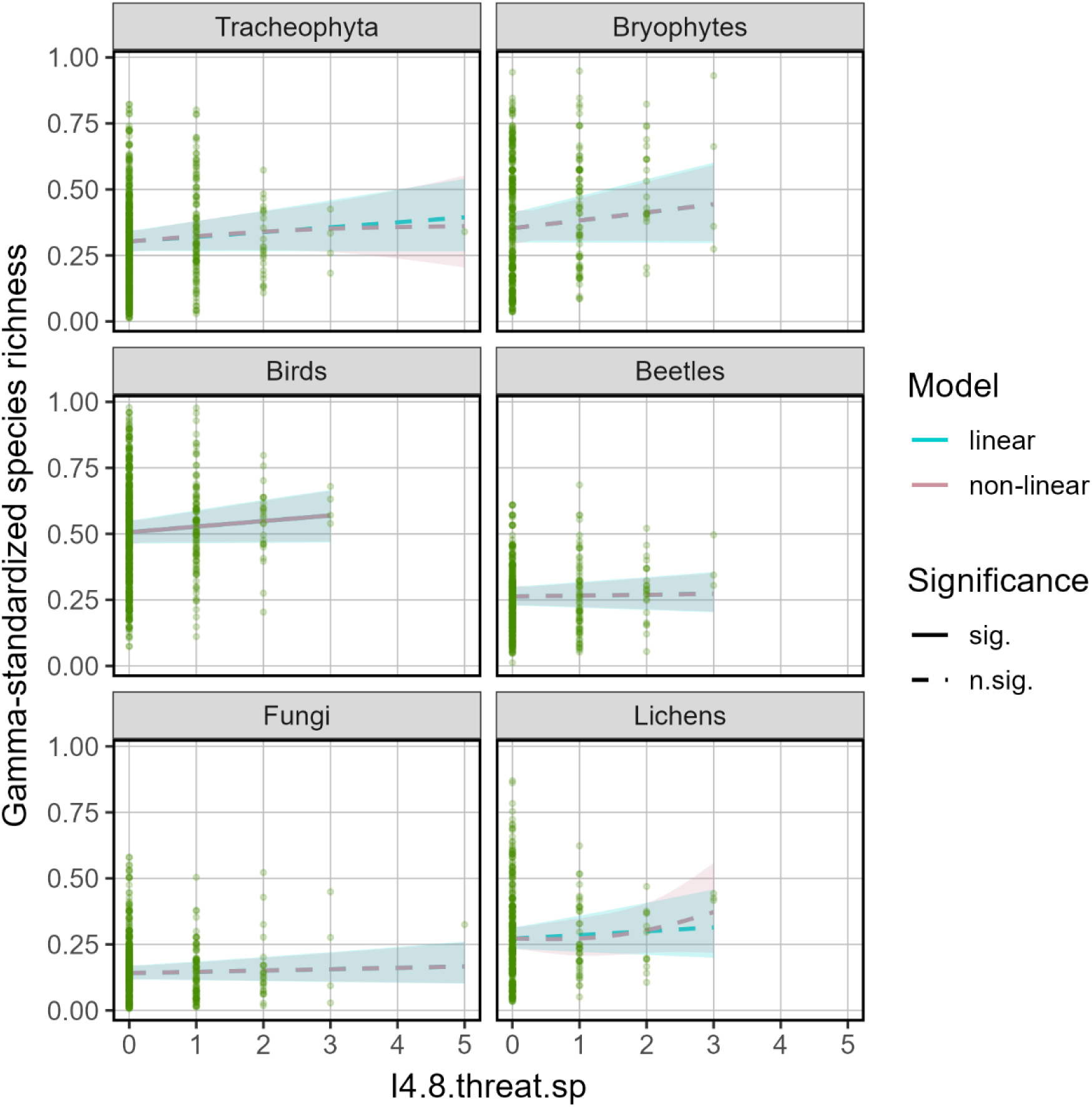

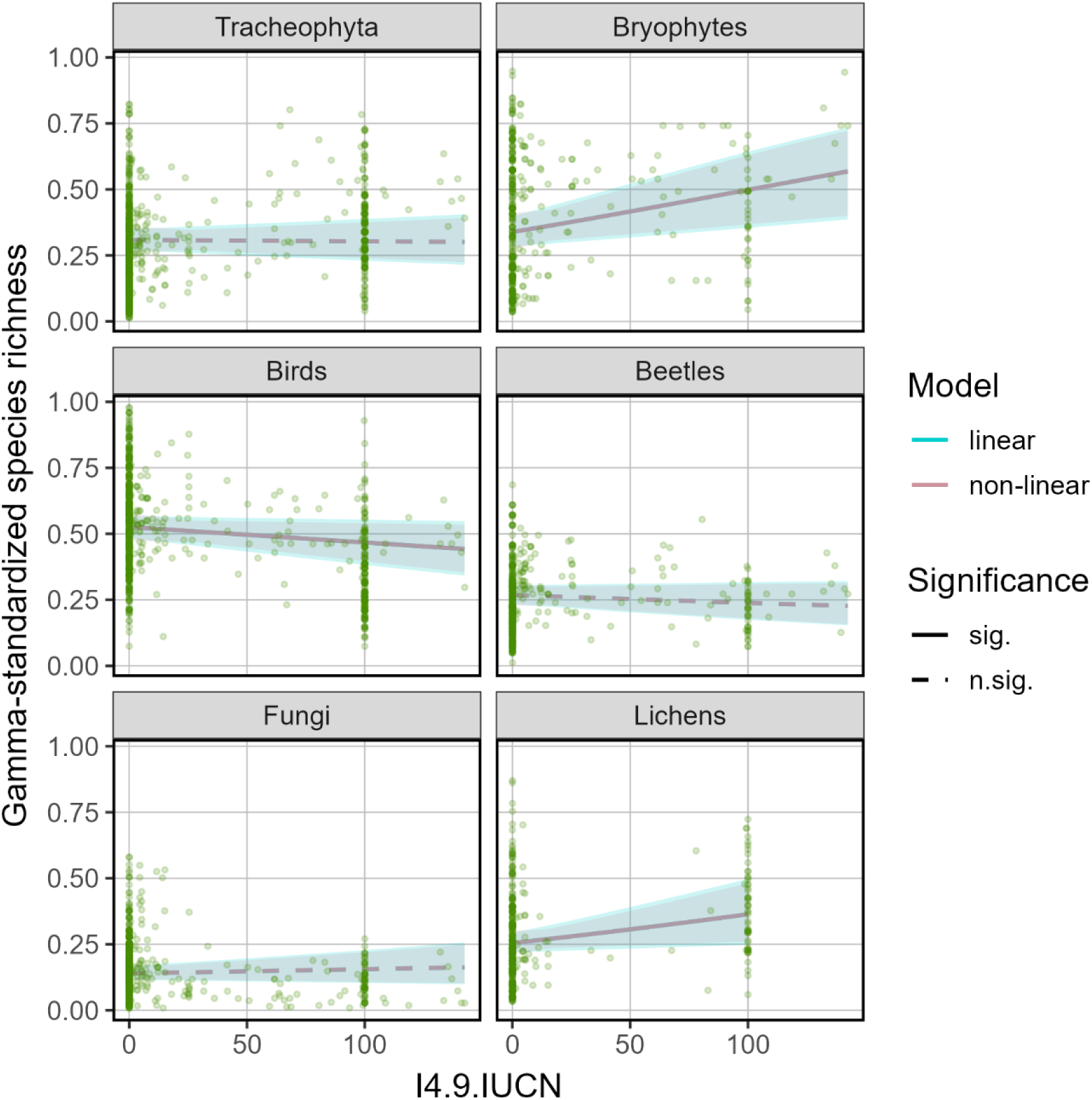

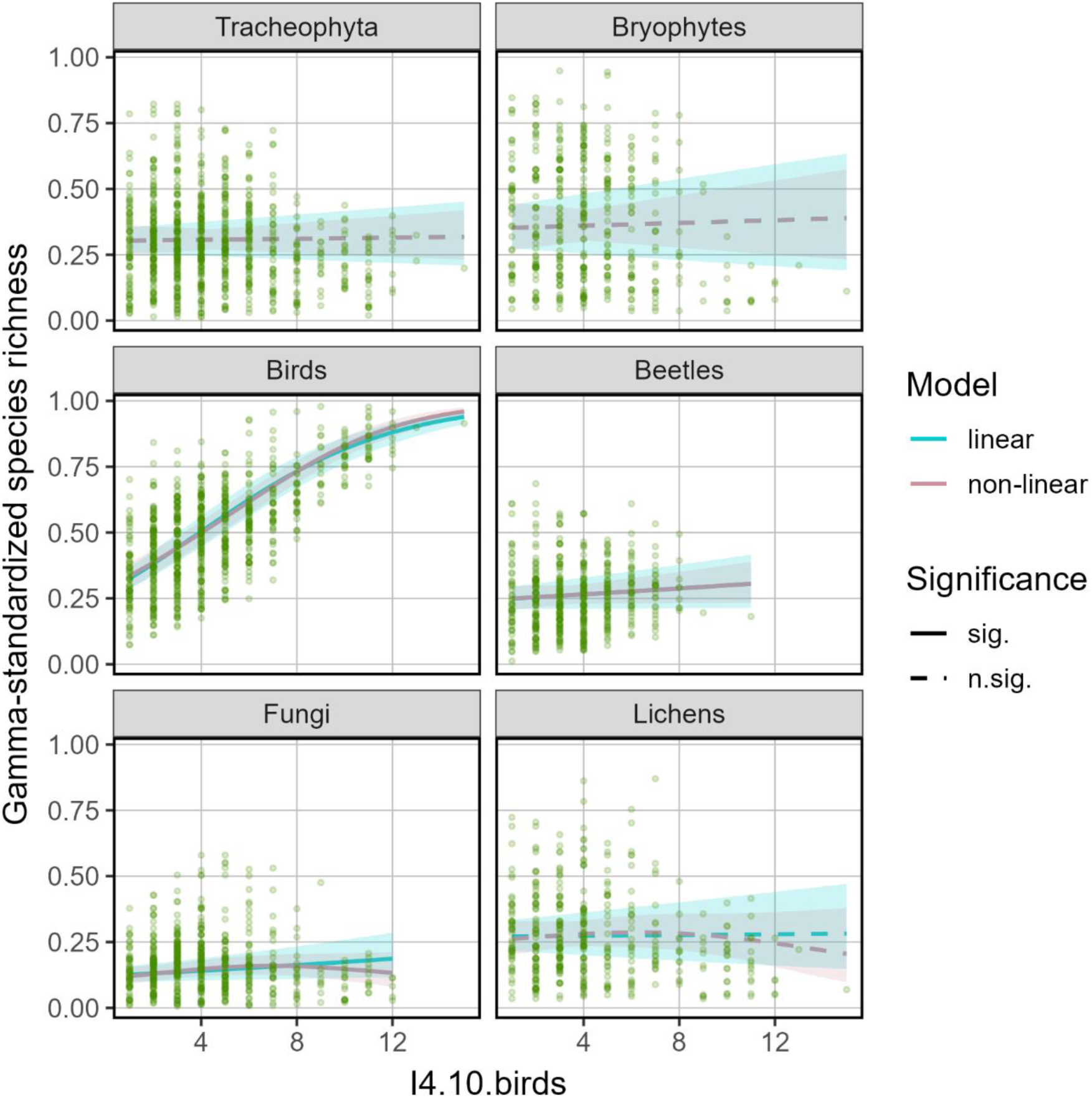

## Appendix 4

Correlation between indicators of Forest Europe from the different taxa datasest. Figures on the right are Pearson correlation coefficients, biplots on the left and histograms on the diagonal represent the distribution of the data. Regeneration (4.2) and Naturalness (4.3) are categorical variables not represented here.

### 3.1. Tracheophytes

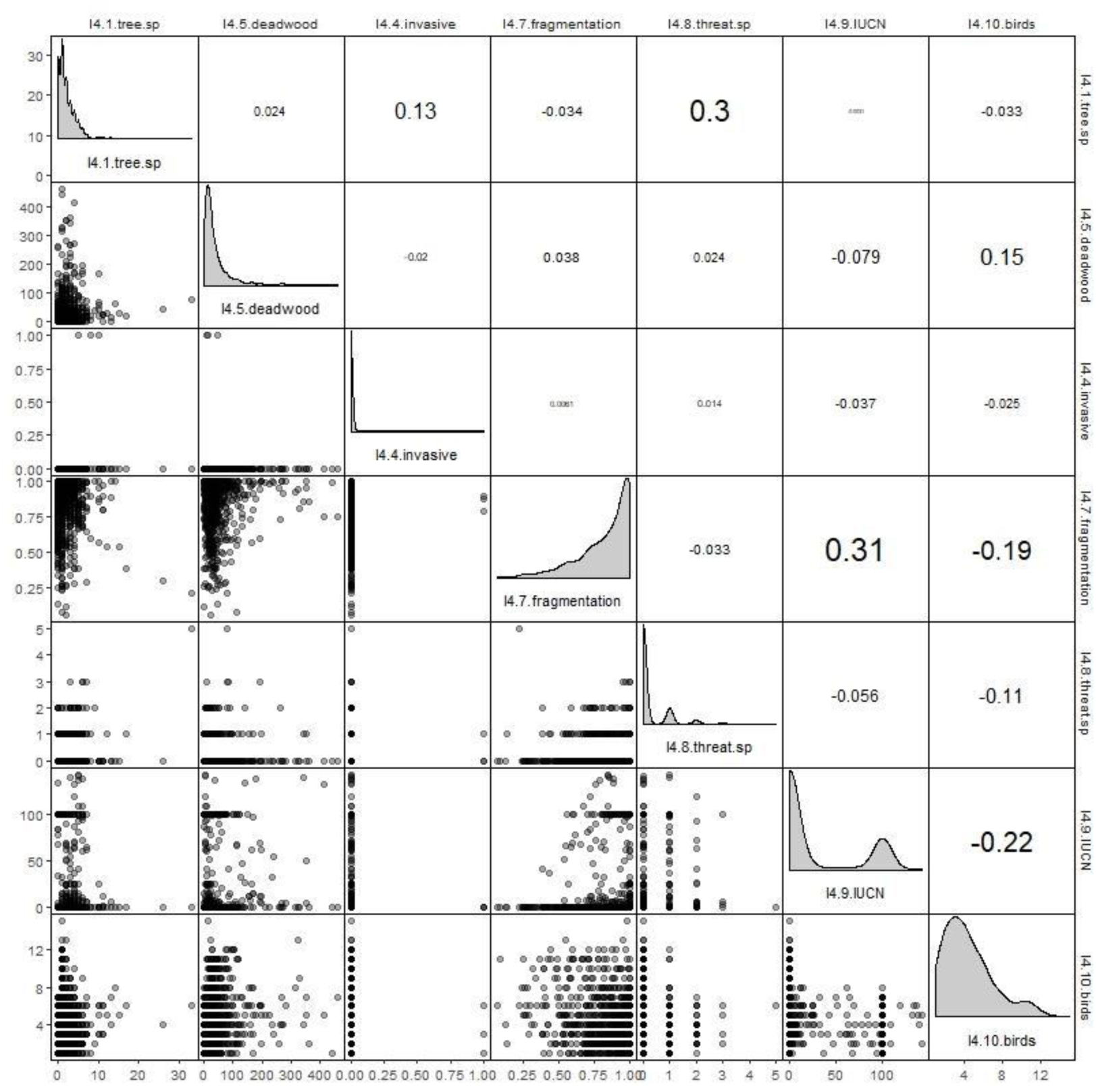

### 3.2. Bryophytes

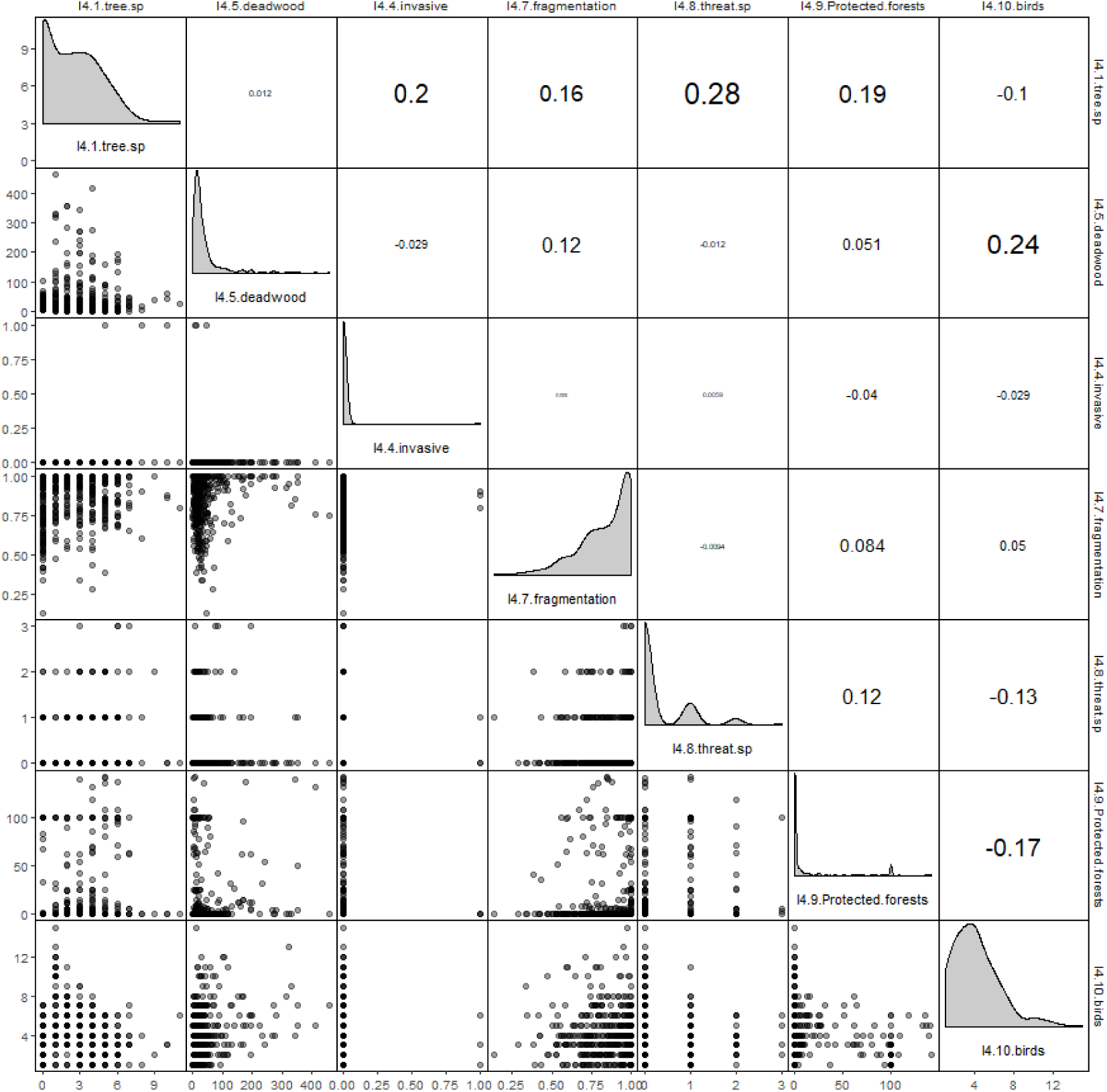

### 3.3. Beetles

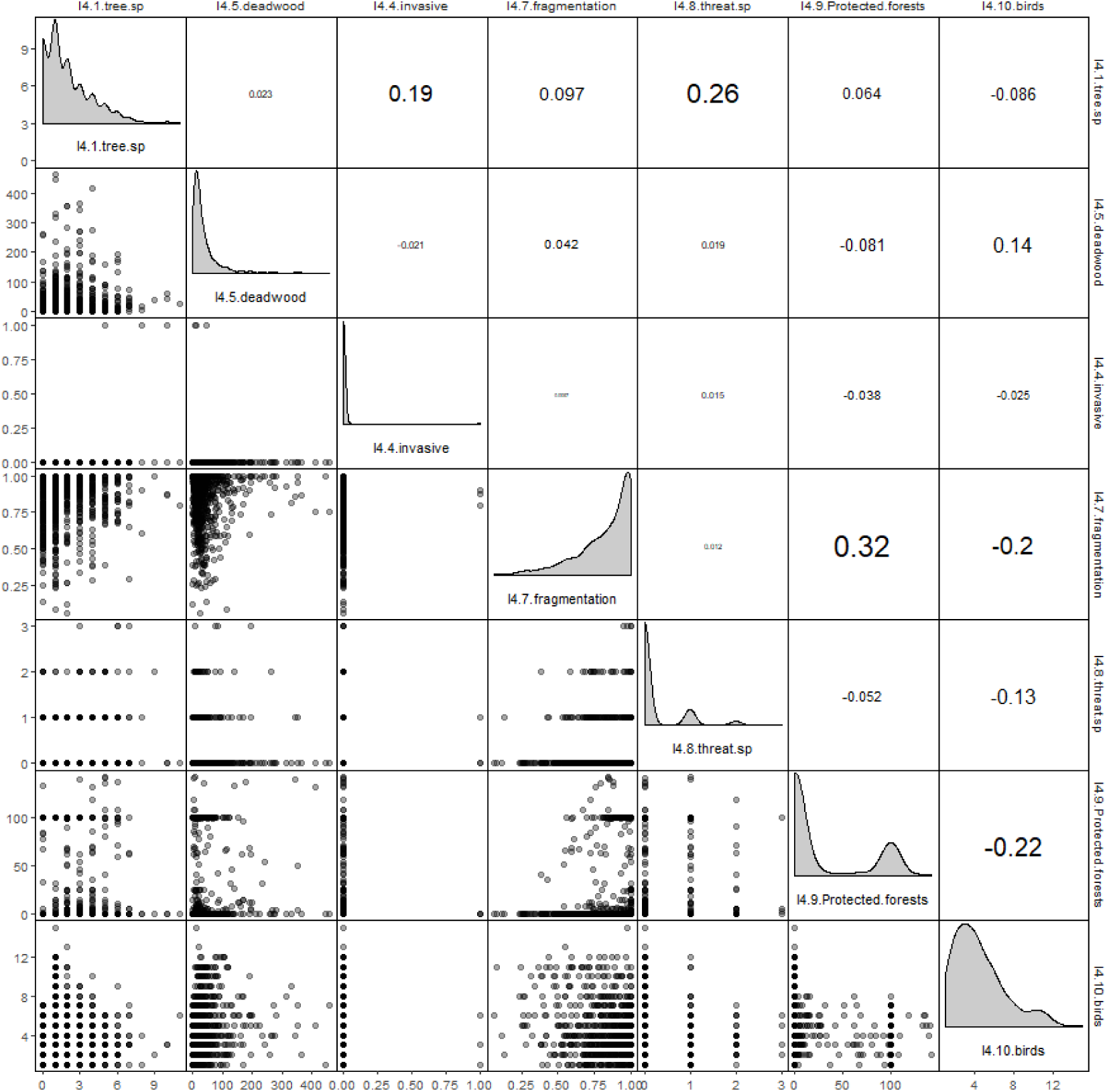

### 3.4. Birds

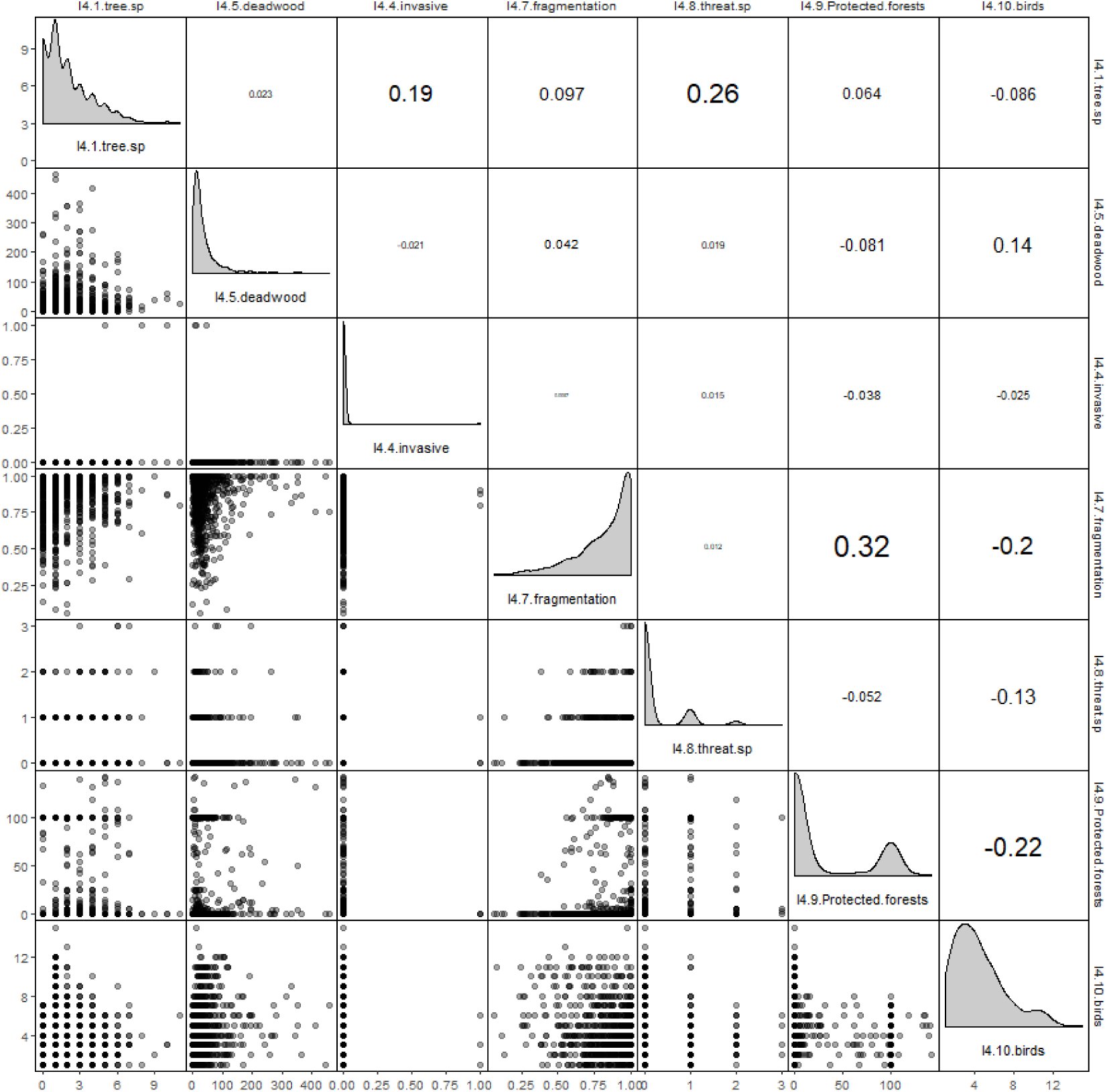

### 3.5. Lichens

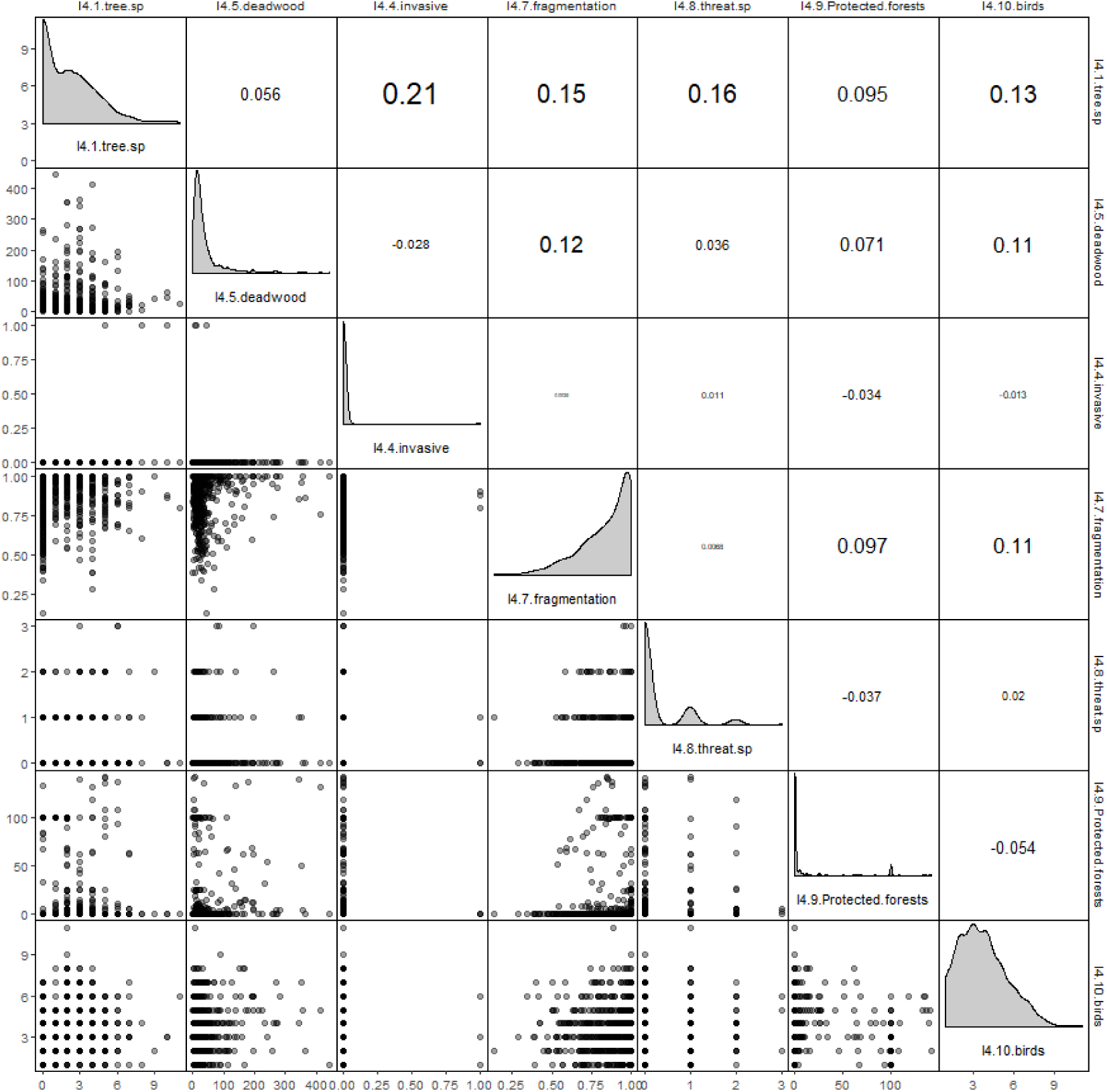

### 3.6. Fungi

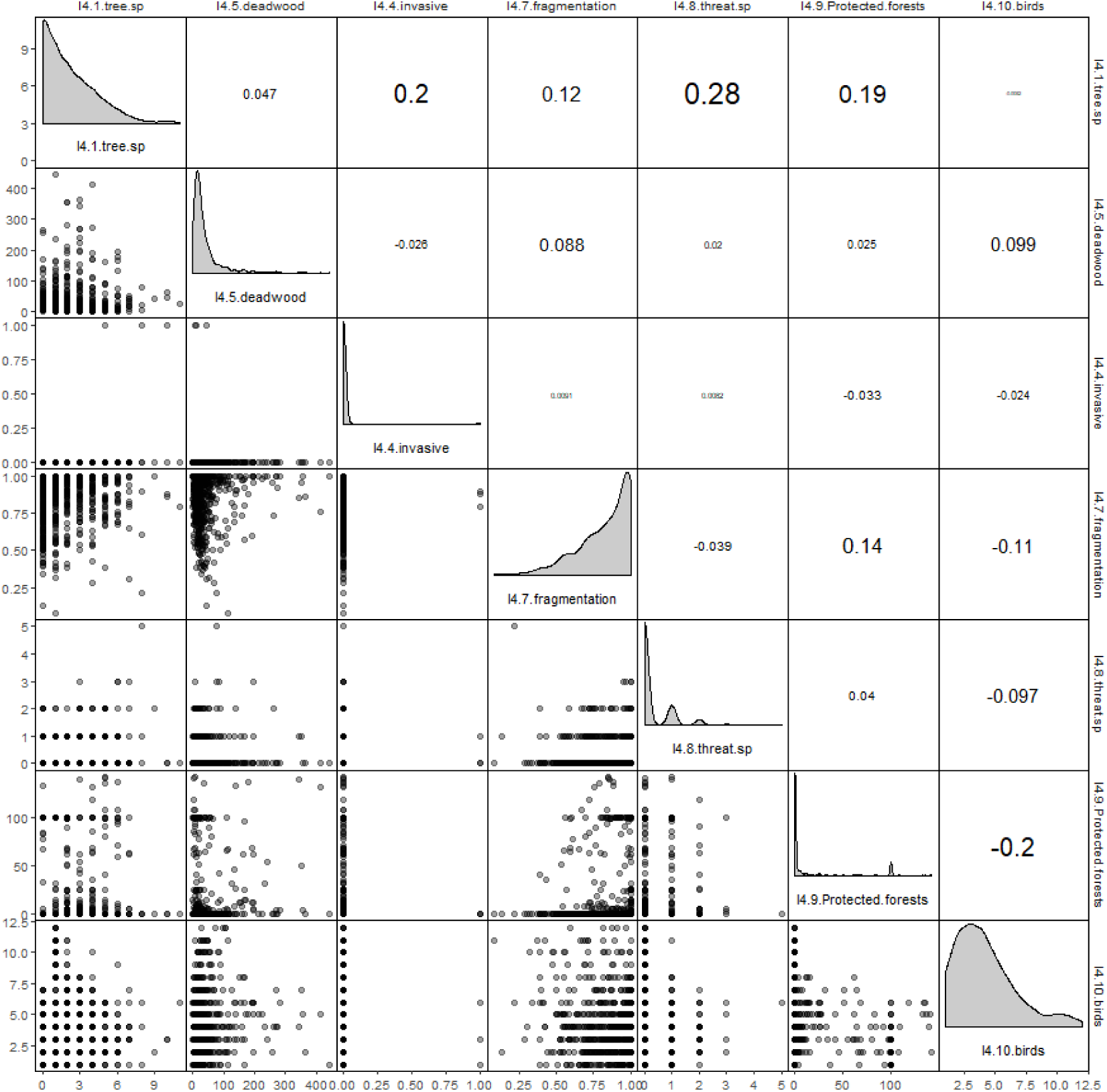

## Appendix 5

Scaled richness estimates table for all single generalized mixed models with beta error distribution and logit link. Intercept for I4.2.Regeneration is “Coppice”, and for I4.3.Naturalness “Plantation” (PLA). S.N = semi-natural forest, UNM = Unmanaged forests. Se = standard error of the mean, pl = critical probability, (*) p<0.1, * p<0.05, ** p<0.01, *** p<0.001.

| Group | variable | estimate | se | p |  |
| --- | --- | --- | --- | --- | --- |
| Tracheophytes | l4.1.tree.sp | 0.0648 | 0.0359 | 0.0712 | (*) |
|  | l4.2.regeneration(natural regeneration) | 0.1431 | 0.2606 | 0.5828 |  |
|  | l4.2.regeneration(planting) | 0.5186 | 0.2754 | 0.0597 | (*) |
|  | l4.3.naturalnessUNM | -0.3878 | 0.0889 | <0.001 | *** |
|  | l4.3.naturalnessS.N | -0.2073 | 0.1429 | 0.147 |  |
|  | l4.4.invasive | 0.0376 | 0.0193 | 0.0508 | (*) |
|  | l4.5.deadwood | 0.0318 | 0.0219 | 0.1463 |  |
|  | l4.7.fragmentation | -0.0898 | 0.0256 | <0.001 | *** |
|  | l4.8.threat.sp | 0.0428 | 0.0221 | 0.0522 | (*) |
|  | l4.9.protected.forests | -0.0087 | 0.0416 | 0.8344 |  |
|  | l4.10.birds | 0.0122 | 0.0326 | 0.707 |  |
| Bryophytes | l4.1.tree.sp | 0.2828 | 0.0633 | <0.001 | *** |
|  | l4.2.regeneration(natural regeneration) | 0.4156 | 0.4546 | 0.3606 |  |
|  | l4.2.regeneration(planting) | -0.5511 | 0.4716 | 0.2426 |  |
|  | l4.3.naturalnessUNM | 0.8606 | 0.128 | <0.001 | *** |
|  | l4.3.naturalnessS.N | 1.5727 | 0.1745 | <0.001 | *** |
|  | l4.4.invasive | -0.0409 | 0.0388 | 0.2917 |  |
|  | l4.5.deadwood | 0.2097 | 0.04 | <0.001 | *** |
|  | l4.7.fragmentation | 0.1384 | 0.0486 | 0.0044 | ** |
|  | l4.8.threat.sp | 0.0776 | 0.0402 | 0.0539 | (*) |
|  | l4.9.protected.forests | 0.2268 | 0.0585 | <0.001 | *** |
|  | l4.10.birds | 0.0235 | 0.0571 | 0.6806 |  |
| Birds | l4.1.tree.sp | 0.0341 | 0.0323 | 0.2916 |  |
|  | l4.2.regeneration(natural regeneration) | 0.0996 | 0.2114 | 0.6375 |  |
|  | l4.2.regeneration(planting) | 0.0452 | 0.2249 | 0.8407 |  |
|  | l4.3.naturalnessUNM | 0.0388 | 0.0785 | 0.6207 |  |
|  | I4.3.naturalnessS.N | 0.2011 | 0.1259 | 0.1104 |  |
|  | I4.4.invasive | -0.0177 | 0.0186 | 0.3417 |  |
|  | I4.5.deadwood | 0.0402 | 0.0203 | 0.0469 | * |
|  | I4.7.fragmentation | -0.1018 | 0.0243 | <0.001 | *** |
|  | I4.8.threat.sp | 0.043 | 0.0205 | 0.0365 | * |
|  | I4.9.protected.forests | -0.1031 | 0.041 | 0.012 | * |
|  | I4.10.birds | 0.6521 | 0.0218 | <0.001 | *** |
| Beetles | I4.1.tree.sp | 0.0242 | 0.034 | 0.4762 |  |
|  | I4.2.regeneration(natural regeneration) | 0.1411 | 0.2247 | 0.53 |  |
|  | I4.2.regeneration(planting) | 0.1782 | 0.233 | 0.4444 |  |
|  | I4.3.naturalnessUNM | -0.0338 | 0.0638 | 0.5965 |  |
|  | I4.3.naturalnessS.N | -0.0749 | 0.109 | 0.4917 |  |
|  | I4.4.invasive | -0.0135 | 0.0175 | 0.441 |  |
|  | I4.5.deadwood | 0.0045 | 0.0204 | 0.8238 |  |
|  | I4.7.fragmentation | 0.0435 | 0.0263 | 0.0983 | (*) |
|  | I4.8.threat.sp | 0.0096 | 0.0196 | 0.6255 |  |
|  | I4.9.protected.forests | -0.0526 | 0.0357 | 0.1409 |  |
|  | I4.10.birds | 0.05 | 0.023 | 0.0297 | * |
| Fungi | I4.1.tree.sp | 0.1476 | 0.0343 | <0.001 | *** |
|  | I4.2.regeneration(natural regeneration) | 0.1607 | 0.2444 | 0.5109 |  |
|  | I4.2.regeneration(planting) | -0.3632 | 0.2542 | 0.1531 |  |
|  | I4.3.naturalnessUNM | 0.5017 | 0.0691 | <0.001 | *** |
|  | I4.3.naturalnessS.N | 0.9083 | 0.1434 | <0.001 | *** |
|  | I4.4.invasive | 0.0035 | 0.0189 | 0.8526 |  |
|  | I4.5.deadwood | 0.2228 | 0.0181 | <0.001 | *** |
|  | I4.7.fragmentation | 0.054 | 0.0292 | 0.0644 | (*) |
|  | I4.8.threat.sp | 0.0226 | 0.022 | 0.3039 |  |
|  | I4.9.protected.forests | 0.0422 | 0.0466 | 0.366 |  |
|  | I4.10.birds | 0.1021 | 0.0324 | 0.0016 | ** |
| Lichens | I4.1.tree.sp | 0.1714 | 0.0507 | <0.001 | *** |
|  | I4.2.regeneration(natural regeneration) | 0.0169 | 0.258 | 0.9479 |  |
|  | l4.2.regeneration(planting) | -0.5447 | 0.2736 | 0.0465 | * |
|  | l4.3.naturalnessUNM | 0.5513 | 0.1028 | <0.001 | *** |
|  | l4.3.naturalnessS.N | 0.7687 | 0.2439 | 0.0016 | ** |
|  | l4.4.invasive | 0.0092 | 0.0313 | 0.7691 |  |
|  | l4.5.deadwood | 0.1139 | 0.0343 | <0.001 | *** |
|  | l4.7.fragmentation | 0.0137 | 0.0442 | 0.7564 |  |
|  | l4.8.threat.sp | 0.0339 | 0.0348 | 0.329 |  |
|  | l4.9.protected.forests | 0.1778 | 0.061 | 0.0036 | ** |
|  | l4.10.birds | 0.0067 | 0.0559 | 0.904 |  |

## References

Alterio, E., Campagnaro, T., Sallustio, L., Burrascano, S., Casella, L., Sitzia, T., 2023. Forest management plans as data source for the assessment of the conservation status of European Union habitat types. Frontiers in Forests and Global Change 5.

Arroyo-Rodríguez, V., Fahrig, L., Tabarelli, M., Watling, J.I., Tischendorf, L., Benchimol, M., Cazetta, E., Faria, D., Leal, I.R., Melo, F.P.L., Morante-Filho, J.C., Santos, B.A., Arasa-Gisbert, R., Arce-Peña, N., Cervantes-López, M.J., Cudney-Valenzuela, S., Galán-Acedo, C., San-José, M., Vieira, I.C.G., Slik, J.W.F., Nowakowski, A.J., Tscharntke, T., 2020. Designing optimal human-modified landscapes for forest biodiversity conservation. Ecology Letters 23, 1404–1420.

Atrena, A., Banelytė, G.G., Bruun, H.H., Goldberg, I., Rahbek, C., Heilmann-Clausen, J., 2024. Plant communities and their relations to habitat and microhabitat features along a management gradient in beech forests in Denmark. Forest Ecology and Management 569.

Barton, K., 2023. MuMIn: multi-model inference. in R package version 1.47.5.

Betts, M.G., Wolf, C., Ripple, W.J., Phalan, B., Millers, K.A., Duarte, A., Butchart, S.H.M., Levi, T., 2017. Global forest loss disproportionately erodes biodiversity in intact landscapes. Nature 547, 441–444.

Boch, S., Prati, D., Hessenmöller, D., Schulze, E.D., Fischer, M., 2013. Richness of Lichen Species, Especially of Threatened Ones, Is Promoted by Management Methods Furthering Stand Continuity. PLoS ONE 8.

Boch, S., Saiz, H., Allan, E., Schall, P., Prati, D., Schulze, E.-D., Hessenmöller, D., Sparrius, L.B., Fischer, M., 2021. Direct and Indirect Effects of Management Intensity and Environmental Factors on the Functional Diversity of Lichens in Central European Forests, In Microorganisms.

Bouget, C., Larrieu, L., Nusillard, B., Parmain, G., 2013. In search of the best local habitat drivers for saproxylic beetle diversity in temperate deciduous forests. Biodiversity and Conservation 22, 2111–2130.

Brooks, M.E., Kristensen, K., van Benthem, K.J., Magnusson, A., Berg, C.W., Nielsen, A., Skaug, H.J., Maechler, M., Bolker, B.M., 2017. glmmTMB Balances Speed and Flexibility Among Packages for Zero-inflated Generalized Linear Mixed Modeling. The R Journal, 9, 378–400.

Burnham, K.P., Anderson, D.R., 2002. Model Selection and Multimodel Inference: A Practical Information-Theoretic Approach (2nd ed). Burrascano, S., Chianucci, F., Trentanovi, G., Kepfer-Rojas, S., Sitzia, T., Tinya, F., Doerfler, I., Paillet, Y., Nagel, T.A., Mitic, B., Morillas, L., Munzi, S., Van der Sluis, T., Alterio, E., Balducci, L., de Andrade, R.B., Bouget, C., Giordani, P., Lachat, T., Matosevic, D., Napoleone, F., Nascimbene, J., Paniccia, C., Roth, N., Aszalós, R., Brazaitis, G., Cutini, A., D’Andrea, E., De Smedt, P., Heilmann-Clausen, J., Janssen, P., Kozák, D., Mårell, A., Mikoláš, M., Nordén, B., Matula, R., Schall, P., Svoboda, M., Ujhazyova, M., Vandekerkhove, K., Wohlwend, M., Xystrakis, F., Aleffi, M., Ammer, C., Archaux, F., Asbeck, T., Avtzis, D., Ayasse, M., Bagella, S., Balestrieri, R., Barbati, A., Basile, M., Bergamini, A., Bertini, G., Biscaccianti, A.B., Boch, S., Bölöni, J., Bombi, P., Boscardin, Y., Brunialti, G., Bruun, H.H., Buscot, F., Byriel, D.B., Campagnaro, T., Campanaro, A., Chauvat, M., Ciach, M., Čiliak, M., Cistrone, L., Pereira, J.M.C., Daniel, R., De Cinti, B., De Filippo, G., Dekoninck, W., Di Salvatore, U., Dumas, Y., Elek, Z., Ferretti, F., Fotakis, D., Frank, T., Frey, J., Giancola, C., Gomoryová, E., Gosselin, M., Gosselin, F., Gossner, M.M., Götmark, F., Haeler, E., Hansen, A.K., Hertzog, L., Hofmeister, J., Hošek, J., Johannsen, V.K., Justensen, M.J., Korboulewsky, N., Kovács, B., Lakatos, F., Landivar, C.M., Lens, L., Lingua, E., Lombardi, F., Máliš, F., Marchino, L., Marozas, V., Matteucci, G., Mattioli, W., Møller, P.F., Müller, J., Németh, C., Ónodi, G., Parisi, F., Perot, T., Perret, S., Persiani, A.M., Portaccio, A., Posillico, M., Preikša, Ž., Rahbek, C., Rappa, N.J., Ravera, S., Romano, A., Samu, F., Scheidegger, C., Schmidt, I.K., Schwegmann, S., Sicuriello, F., Spinu, A.P., Spyroglou, G., Stillhard, J., Topalidou, E., Tøttrup, A.P., Ujházy, K., Veres, K., Verheyen, K., Weisser, W.W., Zapponi, L., Ódor, P., 2023. Where are we now with European forest multi-taxon biodiversity and where can we head to? Biological Conservation 284.

Burrascano, S., De Andrade, R.B., Paillet, Y., Ódor, P., Antonini, G., Bouget, C., Campagnaro, T., Gosselin, F., Janssen, P., Persiani, A.M., Nascimbene, J., Sabatini, F.M., Sitzia, T., Blasi, C., 2018. Congruency across taxa and spatial scales: are we asking too much of species data? Global Ecology and Biogeography 27, 980–990.

Campagnaro, T., Brundu, G., Sitzia, T., 2018. Five major invasive alien tree species in European Union forest habitat types of the Alpine and Continental biogeographical regions. Journal for Nature Conservation 43, 227–238.

Cavard, X., Macdonald, S.E., Bergeron, Y., Chen, H.Y.H., 2011. Importance of mixedwoods for biodiversity conservation: Evidence for understory plants, songbirds, soil fauna, and ectomycorrhizae in northern forests. Environmental Reviews 19, 142–161.

Chaudhary, A., Burivalova, Z., Koh, L.P., Hellweg, S., 2016. Impact of Forest Management on Species Richness: Global Meta-Analysis and Economic Trade-Offs. Scientific Reports 6, 23954.

Chianucci, F., Napoleone, F., Ricotta, C., Ferrara, C., Fusaro, L., Balducci, L., Trentanovi, G., Bradley, O., Kovacs, B., Mina, M., Cerabolini, B.E.L., Vandekerkhove, K., De Smedt, P., Lens, L., Hertzog, L., Verheyen, K., Hofmeister, J., Hošek, J., Matula, R., Doerfler, I., Müller, J., Weisser, W.W., Helback, J., Schall, P., Fischer, M., Heilmann-Clausen, J., Riis-Hansen, R., Goldberg, I., Aude, E., Kepfer-Rojas, S., Kappel Schmidt, I., Riis Nielsen, T., Mårell, A., Dumas, Y., Janssen, P., Paillet, Y., Archaux, F., Xystrakis, F., Tinya, F., Ódor, P., Aszalós, R., Bölöni, J., Cutini, A., Bagella, S., Sitzia, T., Brazaitis, G., Marozas, V., Ujházyová, M., Ujházy, K., Máliš, F., Nordén, B., Burrascano, S., in press. Silvicultural regime shapes understory functional structure in European forests. Journal of Applied Ecology n/a.

Chiarucci, A., Bacaro, G., Filibeck, G., Landi, S., Maccherini, S., Scoppola, A., 2012. Scale dependence of plant species richness in a network of protected areas. Biodiversity and Conservation 21, 503–516.

Chirici, G., McRoberts, R.E., Winter, S., Bertini, R., Bröändli, U.B., Asensio, I.A., Bastrup-Birk, A., Rondeux, J., Barsoum, N., Marchetti, M., 2012. National forest inventory contributions to forest biodiversity monitoring. Forest Science 58, 257–268.

Christensen, M., Hahn, K., Mountford, E.P., Odor, P., Standovar, T., Rozenbergar, D., Diaci, J., Wijdeven, S., Meyer, P., Winter, S., Vrska, T., 2005. Dead wood in European beech (Fagus sylvatica) forest reserves. Forest Ecology and Management 210, 267–282.

Convention on Biological Diversity, 2022. Decision CBD/COP/DEC/15/4, ed. United Nations Environment Programme.

European Commission, 2021. New EU Forest Strategy for 2030, In COM(2021) 572 final. ed. t.D. Communication from the Commission to the European Parliament, the European Economic and Social Commitee and the Commitee of the Regions, Brussels, Belgium. European Commission, Joint Research Centre, Vogt, P., Riitters, K., Caudullo, G., Eckhardt, B., Raši, R., 2019. An approach for pan-European monitoring of forest fragmentation. Publications Office.

European Environment, A., 2020. State of nature in the EU – Results from reporting under the nature directives 2013-2018. Publications Office.

European Environment Agency, 2007. European forest types. Categories and types for sustainable forest management reporting and policy, In EEA Technical report. EEA, Copenhagen, Danemark.

FAO, 2024. The State of the World’s Forests 2024 – Forest-sector innovations towards a more sustainable future. FAO, Rome, Italy.

Fick, S.E., Hijmans, R.J., 2017. WorldClim 2: new 1-km spatial resolution climate surfaces for global land areas. International Journal of Climatology 37, 4302–4315.

Forest Europe, 2015. The updated pan-european indicators for sustainable forest management, pp. 1-4. Ministerial Conference on the Protection of Forests in Europe,, Zvolen, Slovak Republic.

Forest Europe, 2020. State of Europe’s Forests 2020, p. 394. Ministerial Conference on the Protection of Forests in Europe - FOREST EUROPE, Liaison Unit Bratislava.

Gao, T., Nielsen, A.B., Hedblom, M., 2015. Reviewing the strength of evidence of biodiversity indicators for forest ecosystems in Europe. Ecological Indicators 57, 420–434.

Graf, M., Seibold, S., Gossner, M.M., Hagge, J., Weiß, I., Bässler, C., Müller, J., 2022. Coverage based diversity estimates of facultative saproxylic species highlight the importance of deadwood for biodiversity. Forest Ecology and Management 517, 120275.

Group on Earth Observation Biodiversity Observation Network, 2008. GEO BON Concept Document, In GEO-V document 20. Geneva, Switzerland.

Heym, M., Uhl, E., Moshammer, R., Dieler, J., Stimm, K., Pretzsch, H., 2021. Utilising forest inventory data for biodiversity assessment. Ecological Indicators 121, 107196.

Honkanen, M., Roberge, J.-M., Rajasärkkä, A., Mönkkönen, M., 2010. Disentangling the effects of area, energy and habitat heterogeneity on boreal forest bird species richness in protected areas. Global Ecology and Biogeography 19, 61–71.

Hsieh, T.C., Ma, K.H., Chao, A., 2016. iNEXT: an R package for rarefaction and extrapolation of species diversity (Hill numbers). Methods in Ecology and Evolution 7, 1451–1456.

IPBES, 2019. Global assessment report on biodiversity and ecosystem services of the Intergovernmental Science-Policy Platform on Biodiversity and Ecosystem Services (Version 1), eds E.D. Brondizio, S., J.N. Settele, H.T., p. 1144 IPBES secretariat, Bonn, Germany.

Jiguet, F., Devictor, V., Julliard, R., Couvet, D., 2012. French citizens monitoring ordinary birds provide tools for conservation and ecological sciences. Acta Oecologica 44, 58–66.

Kempeneers, P., Sedano, F., Seebach, L., Strobl, P., San-Miguel-Ayanz, J., 2011. Data Fusion of Different Spatial Resolution Remote Sensing Images Applied to Forest-Type Mapping. IEEE Transactions on Geoscience and Remote Sensing 49, 4977–4986.

Langridge, J., Delabye, S., Gilg, O., Paillet, Y., Reyjol, Y., Sordello, R., Touroult, J., Gosselin, F., 2023. Biodiversity responses to forest management abandonment in boreal and temperate forest ecosystems: A meta-analysis reveals an interactive effect of time since abandonment and climate. Biological Conservation 287.

Lassauce, A., Paillet, Y., Jactel, H., Bouget, C., 2011. Deadwood as a surrogate for forest biodiversity: Meta-analysis of correlations between deadwood volume and species richness of saproxylic organisms. Ecological Indicators 11, 1027–1039.

Leidinger, J., Blaschke, M., Ehrhardt, M., Fischer, A., Gossner, M.M., Jung, K., Kienlein, S., Kózak, J., Michler, B., Mosandl, R., Seibold, S., Wehner, K., Weisser, W.W., 2021. Shifting tree species composition affects biodiversity of multiple taxa in Central European forests. Forest Ecology and Management 498, 119552.

Lelli, C., Bruun, H.H., Chiarucci, A., Donati, D., Frascaroli, F., Fritz, Ö., Goldberg, I., Nascimbene, J., Tøttrup, A.P., Rahbek, C., Heilmann-Clausen, J., 2019. Biodiversity response to forest structure and management: Comparing species richness, conservation relevant species and functional diversity as metrics in forest conservation. Forest Ecology and Management 432, 707–717.

Likulunga, L.E., Rivera Pérez, C.A., Schneider, D., Daniel, R., Polle, A., 2021. Tree species composition and soil properties in pure and mixed beech-conifer stands drive soil fungal communities. Forest Ecology and Management 502.

Lindenmayer, D.B., Likens, G.E., 2010. The science and application of ecological monitoring. Biological Conservation 143, 1317–1328.

Linser, S., O’Hara, P., 2019. Guidelines for the Development of a Criteria and Indicator Set for Sustainable Forest Management, In Timber and Forest Discussion Paper. p. 91. United Nations and the Food and Agriculture, Organization of the United Nations, Geneva, Switzerland.

Marshall, E., Wintle, B.A., Southwell, D., Kujala, H., 2020. What are we measuring? A review of metrics used to describe biodiversity in offsets exchanges. Biological Conservation 241, 108250.

McFadden, D., 1979. Quantitative Methods for Analyzing Travel Behaviour of Individuals: Some Recent Developments, In Behaviourial Travel Modelling. eds D.A. Hensher, P.R. Stopher, p. 40. Taylor and Francis Group, London, England.

Müller, J., Boch, S., Prati, D., Socher, S.A., Pommer, U., Hessenmöller, D., Schall, P., Schulze, E.D., Fischer, M., 2019. Effects of forest management on bryophyte species richness in Central European forests. Forest Ecology and Management 432, 850–859.

Müller, J., Brustel, H., Brin, A., Bussler, H., Bouget, C., Obermaier, E., Heidinger, I.M.M., Lachat, T., Förster, B., Horak, J., Procházka, J., Köhler, F., Larrieu, L., Bense, U., Isacsson, G., Zapponi, L., Gossner, M.M., 2015. Increasing temperature may compensate for lower amounts of dead wood in driving richness of saproxylic beetles. Ecography 38, 499–509.

Nakagawa, S., Johnson, P.C.D., Schielzeth, H., 2017. The coefficient of determination R2 and intra-class correlation coefficient from generalized linear mixed-effects models revisited and expanded. Journal of the Royal Society Interface 14.

Noss, R.F., 1990. Indicators for monitoring biodiversity - A hierarchical approach. Conservation Biology 4, 355–364.

Paillet, Y., Archaux, F., du Puy, S., Boulanger, V., Debaive, N., Fuhr, M., Gilg, O., Gosselin, F., Guilbert, E., 2018. The indicator side of tree microhabitats: a multi-taxon approach based on bats, birds and saproxylic beetles. Journal of Applied Ecology 55, 2147–2156.

Paillet, Y., Bergès, L., Hjälten, J., Odor, P., Avon, C., Bernhardt-Romermann, M., Bijlsma, R.J., De Bruyn, L., Fuhr, M., Grandin, U., Kanka, R., Lundin, L., Luque, S., Magura, T., Matesanz, S., Meszaros, I., Sebastia, M.T., Schmidt, W., Standovar, T., Tothmeresz, B., Uotila, A., Valladares, F., Vellak, K., Virtanen, R., 2010. Biodiversity Differences between Managed and Unmanaged Forests: Meta-Analysis of Species Richness in Europe. Conservation Biology 24, 101–112.

Parajuli, R., Markwith, S.H., 2023. Quantity is foremost but quality matters: A global meta- analysis of correlations of dead wood volume and biodiversity in forest ecosystems. Biological Conservation 283.

Pascual, U., Balvanera, P., Anderson, C.B., Chaplin-Kramer, R., Christie, M., González- Jiménez, D., Martin, A., Raymond, C.M., Termansen, M., Vatn, A., Athayde, S., Baptiste, B., Barton, D.N., Jacobs, S., Kelemen, E., Kumar, R., Lazos, E., Mwampamba, T.H., Nakangu, B., O’Farrell, P., Subramanian, S.M., van Noordwijk, M., Ahn, S.E., Amaruzaman, S., Amin, A.M., Arias-Arévalo, P., Arroyo-Robles, G., Cantú-Fernández, M., Castro, A.J., Contreras, V., De Vos, A., Dendoncker, N., Engel, S., Eser, U., Faith, D.P., Filyushkina, A., Ghazi, H., Gómez-Baggethun, E., Gould, R.K., Guibrunet, L., Gundimeda, H., Hahn, T., Harmáčková, Z.V., Hernández-Blanco, M., Horcea-Milcu, A.I., Huambachano, M., Wicher, N.L.H., Aydın, C.İ., Islar, M., Koessler, A.K., Kenter, J.O., Kosmus, M., Lee, H., Leimona, B., Lele, S., Lenzi, D., Lliso, B., Mannetti, L.M., Merçon, J., Monroy-Sais, A.S., Mukherjee, N., Muraca, B., Muradian, R., Murali, R., Nelson, S.H., Nemogá-Soto, G.R., Ngouhouo-Poufoun, J., Niamir, A., Nuesiri, E., Nyumba, T.O., Özkaynak, B., Palomo, I., Pandit, R., Pawłowska-Mainville, A., Porter-Bolland, L., Quaas, M., Rode, J., Rozzi, R., Sachdeva, S., Samakov, A., Schaafsma, M., Sitas, N., Ungar, P., Yiu, E., Yoshida, Y., Zent, E., 2023. Diverse values of nature for sustainability. Nature 620, 813–823.

Penone, C., Allan, E., Soliveres, S., Felipe-Lucia, M.R., Gossner, M.M., Seibold, S., Simons, N.K., Schall, P., van der Plas, F., Manning, P., Manzanedo, R.D., Boch, S., Prati, D., Ammer, C., Bauhus, J., Buscot, F., Ehbrecht, M., Goldmann, K., Jung, K., Müller, J., Müller, J.C., Pena, R., Polle, A., Renner, S.C., Ruess, L., Schönig, I., Schrumpf, M., Solly, E.F., Tschapka, M., Weisser, W.W., Wubet, T., Fischer, M., 2019. Specialisation and diversity of multiple trophic groups are promoted by different forest features. Ecology Letters 22, 170–180.

Pereira, H.M., David Cooper, H., 2006. Towards the global monitoring of biodiversity change. Trends in Ecology & Evolution 21, 123–129.

Proença, V., Martin, L.J., Pereira, H.M., Fernandez, M., McRae, L., Belnap, J., Böhm, M., Brummitt, N., García-Moreno, J., Gregory, R.D., Honrado, J.P., Jürgens, N., Opige, M., Schmeller, D.S., Tiago, P., van Swaay, C.A.M., 2017. Global biodiversity monitoring: From data sources to Essential Biodiversity Variables. Biological Conservation 213, 256–263.

R Core Team, 2023. R: A language and environment for statistical computing. R Foundation for Statistical Computing, Vienna, Austria.

Reise, J., Kukulka, F., Flade, M., Winter, S., 2019. Characterising the richness and diversity of forest bird species using National Forest Inventory data in Germany. Forest Ecology and Management 432, 799–811.

Rigal, S., Dakos, V., Alonso, H., Auniņš, A., Benkő, Z., Brotons, L., Chodkiewicz, T., Chylarecki, P., de Carli, E., del Moral, J.C., Domşa, C., Escandell, V., Fontaine, B., Foppen, R., Gregory, R., Harris, S., Herrando, S., Husby, M., Ieronymidou, C., Jiguet, F., Kennedy, J., Klvaňová, A., Kmecl, P., Kuczyński, L., Kurlavičius, P., Kålås, J.A., Lehikoinen, A., Lindström, Å., Lorrillière, R., Moshøj, C., Nellis, R., Noble, D., Eskildsen, D.P., Paquet, J.-Y., Pélissié, M., Pladevall, C., Portolou, D., Reif, J., Schmid, H., Seaman, B., Szabo, Z.D., Szép, T., Florenzano, G.T., Teufelbauer, N., Trautmann, S., van Turnhout, C., Vermouzek, Z., Vikstrøm, T., Voříšek, P., Weiserbs, A., Devictor, V., 2023. Farmland practices are driving bird population decline across Europe. Proceedings of the National Academy of Sciences 120, e2216573120.

Riva, F., Fahrig, L., 2022. The disproportionately high value of small patches for biodiversity conservation. Conservation Letters 15, e12881.

Sills, J., Gómez-González, S., Ochoa-Hueso, R., Pausas, J.G., 2020. Afforestation falls short as a biodiversity strategy. Science 368, 1439–1439.

Simons, N.K., Felipe-Lucia, M.R., Schall, P., Ammer, C., Bauhus, J., Blüthgen, N., Boch, S., Buscot, F., Fischer, M., Goldmann, K., Gossner, M.M., Hänsel, F., Jung, K., Manning, P., Nauss, T., Oelmann, Y., Pena, R., Polle, A., Renner, S.C., Schloter, M., Schöning, I., Schulze, E.-D., Solly, E.F., Sorkau, E., Stempfhuber, B., Wubet, T., Müller, J., Seibold, S., Weisser, W.W., 2021. National Forest Inventories capture the multifunctionality of managed forests in Germany. Forest Ecosystems 8, 5.

Stevenson, S.L., Watermeyer, K., Caggiano, G., Fulton, E.A., Ferrier, S., Nicholson, E., 2021. Matching biodiversity indicators to policy needs. Conservation Biology 35, 522–532.

Storch, F., Boch, S., Gossner, M.M., Feldhaar, H., Ammer, C., Schall, P., Polle, A., Kroiher, F., Müller, J., Bauhus, J., 2023. Linking structure and species richness to support forest biodiversity monitoring at large scales. Annals of Forest Science 80, 3.

Tews, J., Brose, U., Grimm, V., Tielbörger, K., Wichmann, M.C., Schwager, M., Jeltsch, F., 2004. Animal species diversity driven by habitat heterogeneity/diversity: The importance of keystone structures. Journal of Biogeography 31, 79–92.

Tomppo, E., Gschwantner, T., Lawrence, M., Mc Roberts, R.E., 2010. National forest inventories. Pathways for common reporting. Springer Science, Heidelberg, Allemagne.

Trentanovi, G., Campagnaro, T., Sitzia, T., Chianucci, F., Vacchiano, G., Ammer, C., Ciach, M., Nagel, T.A., del Río, M., Paillet, Y., Munzi, S., Vandekerkhove, K., Bravo-Oviedo, A., Cutini, A., D’Andrea, E., De Smedt, P., Doerfler, I., Fotakis, D., Heilmann-Clausen, J., Hofmeister, J., Hošek, J., Janssen, P., Kepfer-Rojas, S., Korboulewsky, N., Kovács, B., Kozák, D., Lachat, T., Mårell, A., Matula, R., Mikoláš, M., Nordén, B., Ódor, P., Perović, M., Pötzelsberger, E., Schall, P., Svoboda, M., Tinya, F., Ujházyová, M., Burrascano, S., 2023. Words apart: Standardizing forestry terms and definitions across European biodiversity studies. Forest Ecosystems 10, 100128.

Weber, D., Hintermann, U., Zangger, A., 2004. Scale and trends in species richness: considerations for monitoring biological diversity for political purposes. Global Ecology and Biogeography 13, 97–104.

Westgate, M.J., Tulloch, A.I.T., Barton, P.S., Pierson, J.C., Lindenmayer, D.B., 2017. Optimal taxonomic groups for biodiversity assessment: a meta-analytic approach. Ecography 40, 539–548.

Wickham, H., 2016. ggplot2: Elegant Graphics for Data Analysis, 2 edn. Springer Cham.

Wildermuth, B., Hagge, J., Seifert, C.L., Tjaden, R., Schuldt, A., 2024. Beneficial effects of native broadleaved forests on canopy beetle diversity are not reduced by admixture of non- native conifers. Journal of Applied Ecology 61, 1000–1014.

Wood, S., 2023. Mixed GAM Computation Vehicle with Automatic Smoothness Estimation. V1.9.

Zeller, L., Baumann, C., Gonin, P., Heidrich, L., Keye, C., Konrad, F., Larrieu, L., Meyer, P., Sennhenn-Reulen, H., Müller, J., Schall, P., Ammer, C., 2022. Index of biodiversity potential (IBP) versus direct species monitoring in temperate forests. Ecological Indicators 136, 108692.

Zeller, L., Förster, A., Keye, C., Meyer, P., Roschak, C., Ammer, C., 2023. What does literature tell us about the relationship between forest structural attributes and species richness in temperate forests? – A review. Ecological Indicators 153, 110383.

Zuur, A.F., Ieno, E.N., Elphick, C.S., 2010. A protocol for data exploration to avoid common statistical problems. Methods in Ecology and Evolution 1, 3–14.

